# The role of two major nucleoid associated proteins in Streptomyces, HupA and HupS, in stress survival and gene expression regulation

**DOI:** 10.1101/2024.06.24.600410

**Authors:** Agnieszka Strzałka, Jakub Mikołajczyk, Klaudia Kowalska, Michał Skurczyński, Neil Holmes, Dagmara Jakimowicz

**Affiliations:** Molecular Microbiology Department, Faculty of Biotechnology, University of Wroclaw, Wroclaw, Poland; The John Innes Centre, Norwich Research Park, Norwich, UK

## Abstract

*Streptomyces* are sporulating soil bacteria with enormous potential for secondary metabolites biosynthesis. Regulatory networks governing *Streptomyces coelicolor* differentiation and secondary metabolites production are complex and composed of numerous regulatory proteins ranging from specific transcriptional regulators to sigma factors. Nucleoid associated proteins (NAPs) are also believed to contribute to regulation of gene expression. Upon DNA binding these proteins impact DNA accessibility. Among NAPs HU proteins are the most widespread and abundant. Unlike other bacteria, the *Streptomyces* genome encodes two HU homologs: HupA and HupS, differing in structure and expression profile. In this study, we explore whether HupA and HupS affect *S. coelicolor* growth under optimal and stressful conditions and how they control global gene expression. By testing both single and double mutants we address the question of both HU homologs complementarity. The lack of both *hup* genes led to growth and sporulation inhibition, as well as increased spore fragility. Our data indicate a synergy between the functions of HupA and HupS during *S. coelicolor* growth. We also demonstrate, that both HU homologs can be considered global transcription regulators influencing expression of numerous genes encoding proteins linked to chromosome topology, secondary metabolites production and transcription. We identify the independent HupA and HupS regulons as well as genes under the control of both HupA and HupS proteins. Our data indicate some extent of redundancy as well as independent function of both homologs.

**Importance:** *Streptomyces* belong to the bacterial family widely used in the production of antibiotics as well as research for new bioactive substances with antimicrobial properties. Gene expression in *Streptomyces*, and consequently the production of secondary metabolites, is controlled by a vast and complex network of transcriptional regulators. Our data indicate that two proteins, HupA and HupS, involved in the maintenance of chromosome structure, also participate in this regulatory network. Their presence appears to important for *S. coelicolor*’s adaptation for survival in unfavorable conditions such as high temperature. The lack of one or both HU proteins affects the expression of many genes, indicating that they act as global transcriptional regulators.

## Introduction

Nucleoid associated proteins (NAPs) are bacterial proteins that perform a role similar to eukaryotic histones. By coating, bridging and bending DNA molecules, these proteins organize and compact DNA. They also share other properties with histones, such as small size, a high content of basic amino acids, and a lack of (or very low) DNA sequence specificity. What is more, similarly to histones, by modifying the accessibility of DNA to transcriptional machineries, they play a role in the regulation of gene expression (1, 2).

In many bacterial cells, the most abundant NAP is a small, positively charged protein HU (Heat Unstable) (3). In *Escherichia coli,* HU abundance, estimated to be between 30, 000 and 55, 000 HU molecules per cell, reaches its peak during exponential growth (4). Like other NAPs, this protein exhibits little sequence specificity, but shows increased affinity to supercoiled, single stranded or distorted DNA (5–7). Interestingly, the impact of HU binding on DNA structure *in vitro* depends on protein/DNA molar ratio and osmolarity. At law salt concentration, HU promotes DNA compaction, while at high salt concentration, its binding leads to the formation of “rigid filaments” (8–10). *In vivo* HU homologs affect gene expression and change the distribution of RNA polymerase by altering DNA topology within promoter regions or promoting long-range DNA contacts (11, 12). Recent studies have revealed that HU homologs influence transcription of genes connected to stress response or virulence in many bacterial species including: *E. coli* (*13, 14*) *Salmonella enterica* (15), *Vibrio parahaemolyticus* (*16*), *Francisella tularensis* (17) and *Helicobacter pylori* (18).

Most of the Proteobacteria and Bacteroidetes, like *E. coli*, possess two HU homologs that form homo- or heterodimers, while other phyla have only one HU homolog which forms a homodimer (19). In *E. coli* two HU homologs, HUα and HUβ, share 69% amino acid identity (20), but differ in affinity for different DNA structures, binding modes and their level during culture growth (5, 21, 22). Notably, actinobacterial HU homologs form a distinct group and are characterized by the presence of a long, positively charged C-terminal domain (23). In *Mycobacteria,* the presence of C-terminal domain is necessary for DNA binding (24, 25). Interestingly, a few actinobacterial orders, namely *Streptomycetales, Propionibacteriales, Kineosporiales and Micrococcales,* possess two HU homologs differing in structure: one with an extended C-terminal domain and one similar to *E. coli* HU.

*Streptomyces* are soil-dwelling bacteria known for antibiotic production and a complex life cycle that includes sporulation. *Streptomyces coelicolor* long, linear chromosome (8.6 Mbp) contains more than 20 biosynthetic gene clusters involved in production of secondary metabolites (26). Most of them remain inactive during growth, while a few are activated before the start of sporulation (27). The structure of the *Streptomyces* chromosome also changes during growth, from uncondensed in vegetative hyphae to tightly packaged in unigenomic spores or late stationary phase (28, 29). These changes of *Streptomyces* chromosome organisation seem to be related, among other factors, to changes in the levels of two HU homologs. According to proteomic and transcriptomic data, HupA is the most abundant NAP during vegetative growth (30, 31) and binds preferentially at the central region of chromosome (32). HupS levels increase during sporulation and the protein shows enhanced binding in terminal regions of chromosome (29). In contrast to *E. coli* HU homologs, HupA and HupS share little sequence homology. HupA is more similar to the canonical *E. coli* HU, but has only 38% identity with N-terminal domain of HupS. The long positively charged C-terminal domain of HupS contains multiple lysine repeats, similar to those found in other Actinobacterial proteins such as topoisomerase I (TopA) (30, 33–35). Interestingly, the lack of only one HU homolog moderately affects *Streptomyces* growth. Deletion of *hupA* in both *S. coelicolor* and *S. lividans* reduced their growth rate (32, 36), while *hupS* deletion in *S. coelicolor* and *S. venezuelae* resulted in diminished compaction of chromosome in spores, which were more sensitive to high temperatures than wild type spores (29, 33).

Given the minimal phenotype of either *hupA* or *hupS* deletion mutants, we expected that the functions of HupA and HupS may be partially complementary. The cooperation between HupA and HupS has not yet been described. Thus, in this work we sought to examine the consequences of *hupA* and *hupS* deletion for *S. coelicolor* growth under optimal and stressful conditions. Given that NAPs binding impacts transcription, the phenotype of *hupS* and *hupA* mutant strains could be at least partially attributed to their influence on transcription. Therefore, we also set out to establish HupA and HupS regulons in *S. coelicolor*. We show that both HU homologs regulate genes involved in secondary metabolism and stress response. Comparison of HupA and HupS regulons allowed us to determine the extent of HupA and HupS cooperation. Taken together, our results suggest that, apart from their structural roles, HupA and HupS binding has as impact on global gene expression, facilitating survival under various environmental conditions.

## Materials and Methods

### Growth conditions and genetic modifications of bacterial strains

The *E. coli* and *S. coelicolor* strains used are listed in Supplementary Table S1. The culture conditions, antibiotic concentrations, and transformation and conjugation methods followed the general procedures for *E. coli* (*37*) and *Streptomyces* (*38*). For plate cultures of *S. coelicolor* strains, minimal medium supplemented with 1% mannitol (MM) or soy flour medium (SFM) was used. For the growth rate evaluation, *S. coelicolor* cultures in YEME/TSB were inoculated with spores to final 0.01 U/ml (1 U of spores increases medium absorbance by 1) and cultured in microplates (250 μl per well), for 72h at 30°C using a Bioscreen C (Automated Growth Curves Analysis System, Growth Curves USA), with five experimental replicates for each strain. *S. coelicolor* growth curves were analyzed using the log-logistic model, for each strain half-time was determined.

In order to construct strain lacking *hupA* and *hupS* genes we plasmid pKF289 (33) was introduced into ASMK011 strain (*ΔhupA::scar*). After conjugation colonies resistant to hygromycin were obtained yielding strain ASMK019 (*ΔhupA::scar ΔhupS::higro*), which was verified using PCR. In order to create a complementation strain, *hupA* gene with its native promoter was amplified using *hupA_pSET_FW* and *hupA_pSET_RV* primers, and then ligated into a pSET152 vector. Obtained was plasmid was introduced into ASMK019 strain. After conjugation colonies resistant to hygromycin and apramycin were obtained yielding strain ASMK019.2 (*ΔhupA::scar ΔhupS::higro* pSET152*hupA*). DNA manipulations were carried out by standard protocols (37). The genetic modifications of the obtained strains were verified by PCR and sequencing. The oligonucleotides used for PCR are listed in Supplementary Table S2.

### Stress sensitivity analyses

First, spore concentrations were measured using a Thoma Cell Counting Chamber and Leica DM6 B fluorescence microscope equipped with a 40x objective. Spores were subjected to either increased temperature (60°C for 15-45 min) or detergent (2.5-10% SDS for 1h in room temperature). Next serial dilutions of spores were plated on SFM medium. To test UV sensitivity, spores were first plated and then exposed to UV light for 15-45s. For oxidative stress analysis serial dilutions of spores were plated on the SFM medium containing increasing concentration of H_2_O_2_ (0 – 1 mM) and incubated in 30°C for 5 days. After 5 days of incubation at 30°C the number of growing colonies was counted to determine the percentage of plated spores that survived the stress.

### Microscopy analysis

For microscopy analysis *S. coelicolor* spores were cultured for 44 hours on microscopy coverslips inserted at a 45° angle in a MM solid medium containing 1% mannitol and then mycelia were fixed with a 2.8% paraformaldehyde/0.00875% glutaraldehyde mixture for 10 min at room temperature. After fixation samples were digested with lysozyme (2 mg/ml in 20 mM Tris–HCl supplemented with 10 mM EDTA and 0.9% glucose) for 2 min, washed with PBS, blocked with 2% BSA in a PBS buffer for 10 min and incubated with 0.1–1 μg/ml DAPI (4’, 6-diamidyno-2-fenyloindol, Molecular Probes) and WGA-Texas Red (Wheat Germ Agglutinin-Texas Red) for 60 min. Fluorescence microscopy was performed using a Leica DM6 B fluorescence microscope equipped with a 100x oil immersion objective. Sporulating hyphae were analyzed using custom protocols involving *Fiji* (*39*) and *R* software (40), code is available at https://github.com/astrzalka/sporecounter.

### RNA isolation and RNA-seq bioinformatic analysis

For RNA-seq, total RNA was isolated from 30 ml YEME/TSB cultures. Cultures were inoculated with *S. coelicolo*r spores (amount of spores was normalized by OD measurement of preliminary cultures), and cultivated in flasks with spring coils for 24-36h in 30°C. For osmotic stress experiment growth medium was supplemented with NaCl to final concentration 0.5M. Mycelia were collected at two time points (exponential and early stationary growth, determined individually for each strain based on the growth curve) by collecting 2 ml from the culture and centrifugation. Cell pellets were frozen and stored at -70°C for subsequent RNA isolation. RNA was isolated using the RNeasy Mini Kit (Qiagen) following manufacturer’s instructions, DNA digestions was performed using on column digestion with DNase-I (Qiagen) and TURBO DNase I (Ambion). RNA quality and concentration was measured using Nanodrop (Thermo Fisher Scientific) and Qubit (Thermo Fisher Scientific). The absence of DNA in the sample was confirmed using PCR.

Library preparation and RNA-sequencing was performed by Genewiz (Germany). Trimmomatic software (version 0.39) (41) was used to remove adapter sequences from sequenced reads. Obtained reads were mapped to *S. coelicolor* genome (NC_003888.3) using *Bowtie2* software (version 2.3.5.1) (42, 43) and processed with *samtools* (version 1.10) (44). On average 4 x 10^6^ reads mapped successfully to *S. coelicolor* genome. Differential analysis was performed using R packages *Rsubread* (version 2.10) and *edgeR* (version 3.38) (45, 46) following a protocol described in (47). Gene count matrix was normalized using *egdeR* and quasi-likelihood negative binomial was fitted to the data. Differential expression was tested using *glmTtreat* function with 1.5 fold change threshold. Genes were considered to be differentially expressed if false discovery rate (FDR) was below 0.05 threshold and Log_2_FC value was above 1.5. For data visualization *ggplot2* (version 3.3.6) (48), *ggVennDiagram* (version 1.2.0) (49) and *tidyHeatmap* (version 1.6.0) (50) R packages were used. Cluster analysis was performed using *clust* programme (version 1.10.8) (51). RNA isolation and Reverse-Transcription and Quantitative PCR (RT-qPCR)

RNA for RT-qPCRwas isolated from 5 mL YEME/TSB liquid medium *S. coelicolor* cultures cultivated for 24-36 h. Mycelia were collected by centrifugation, frozen and stored at -70°C for subsequent RNA isolation. RNA was isolated using the RNeasy Mini Kit (Qiagen) following manufacturer’s instructions, digested with TURBO DNase I (Invitrogen) and checked for chromosomal DNA contamination using PCR. A total of 500 ng of RNA was used for cDNA synthesis using the Maxima First Strand cDNA synthesis kit (Thermo Fisher Scientific) in a final volume of 20 μl. Obtained cDNA was diluted to 100 μl and directly used for quantitative PCRs performed with PowerUp SYBR Green Master Mix (Applied Biosystems). The relative level of transcript of interest was quantified using the comparative ΔΔCt method using the *hrdB* transcript as the endogenous control (StepOne Plus real-time PCR system, Applied Biosystems).

## Results and Discussion

### Deletion of *hupA* and *hupS* has a synergistic effect on *Streptomyces* growth and development

Previous reports concerning the role of HU homologs in *Streptomyces* showed only a moderate phenotype of deletion strains; specifically: the Δ*hupA* mutant exhibited slower growth (32, 36), while the Δ*hupS* mutant displayed decreased nucleoid compaction and increased spore sensitivity to thermal stress (29, 33). Given the high level of both proteins in the cell, we wondered if the functions of HU homologs could be redundant in *Streptomyces* and whether HupA or HupS would compensate for the loss of the other homolog. To test this hypothesis, we constructed a double deletion Δ*hupA*Δ*hupS S. coelicolor* strain and compared its growth under various conditions to that of the Δ*hupA* and Δ*hupS* strains.

The growth of Δ*hupA*Δ*hupS* strain was more inhibited than that of either of single mutant (Fig. 1 A). Both the Δ*hupA*Δ*hupS* and Δ*hupA* strains showed a significant delay in the initiation of growth in liquid medium (half time 28.7 and 27.3 h, based on a log-logistic growth model, respectively) compared to wildtype and Δ*hupS* strains (17.5 and 16.3 hours, respectively). However, after the complementation of the double deletion strain with *hupA* gene, an improvement in growth was observed (Fig. S1 A). A similar growth delay was also observed for Δ*hupA*Δ*hupS* strain during culture on solid medium (Fig. 1 B). Moreover, Δ*hupA*Δ*hupS* colonies remained white, as this strain did not produce the characteristic for *S. coelicolor* spores grey-brown pigment (Fig. 1 B). Microscopic observations confirmed, however, that the double deletion strain was able to sporulate (Fig. S1 B), but only after a prolonged incubation (∼7 days) compared to 3 days for the wild type and Δ*hupS* strains, and 4 days for the Δ*hupA* strain. The analysis of sporogenic hyphae in the mutant strains also showed that the Δ*hupA*Δ*hupS* strain had chromosome segregation defects, with 10.4% of spores lacking DNA; in comparison, the wild type strain had only 1.5% anucleate spores. Segregation defects were detectable also in single deletion strains: 3.9% spores lacked DNA in the Δ*hupA* strain and 4.4% in the Δ*hupS* strain (Fig. 1 C, D).

**Figure 1.**
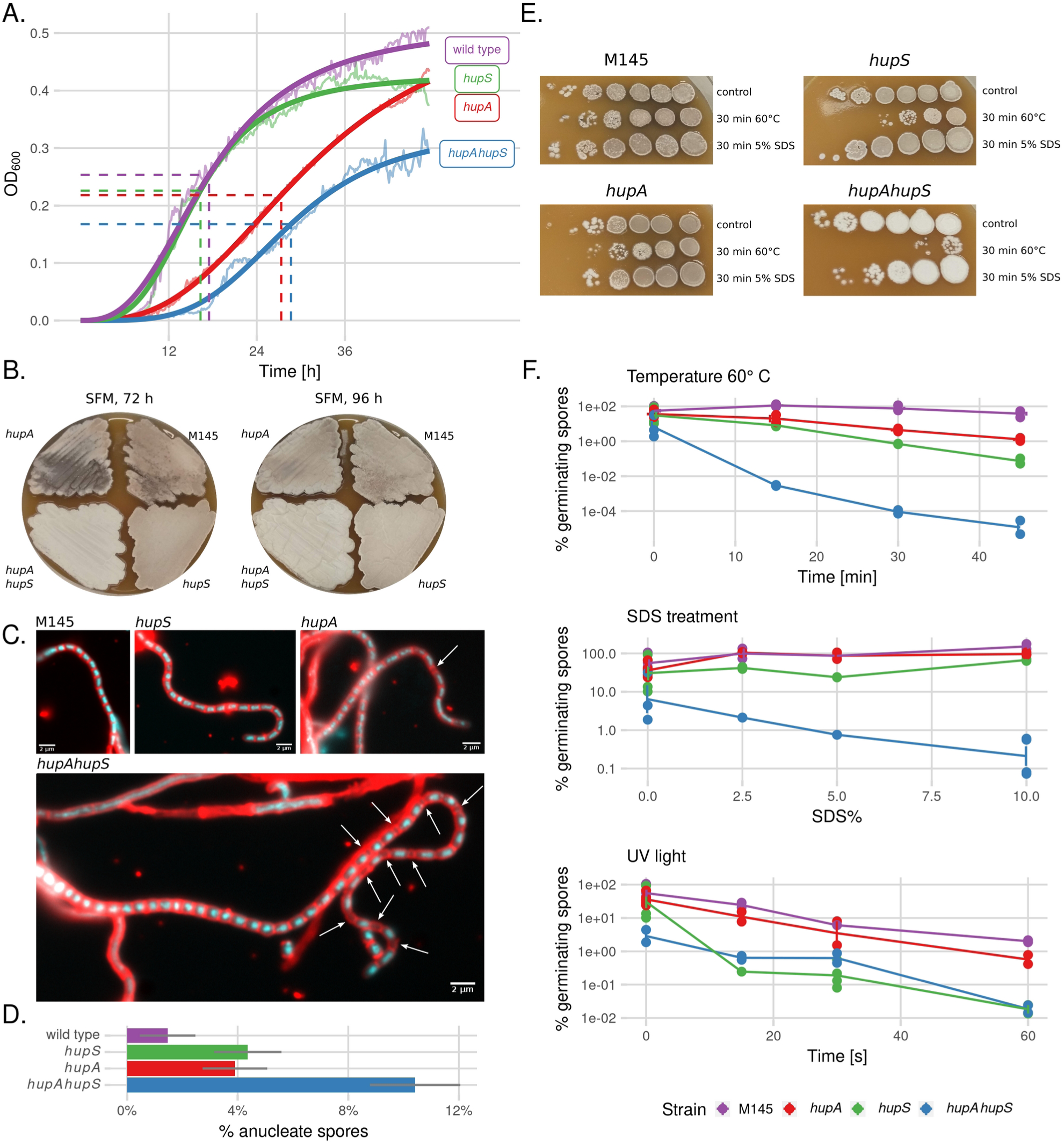
Deletion *hupA* or/and *hupS* inhibits growth and increases stress sensitivity of *S. coelicolor* spores. **A.** Growth curves of wild type (purple), Δ*hupA* (red*), ΔhupS* (green) and Δ*hupAΔhupS* (blue) *s*trains in liquid ‘79’ medium. Bold lines represent log-logistic model while striped lines show half time calculated by the model. **B.** Colonies of wild type, Δ*hupA,* Δ*hupS* and Δ*hupA*Δ*hupS s*trains on solid SFM medium after incubation in 30°C for 72 and 96 hours. Grey appearance of colonies indicates sporulation. **C.** Microscopic images of wild type, Δ*hupA, ΔhupS* and Δ*hupA*Δ*hupS* hyphae stained with DAPI (blue, DNA) and WGA-Texas Red (red, cell wall). White arrows indicate positions of anucleate prepsore compartments. Scale bar 2 μm. **D.** Percentage of anucleate spores in sporulating hyphae of wild type (purple), Δ*hupA* (red), Δ*hupS* (green) and Δ*hupA*Δ*hupS* (blue) *s*trains. Grey lines represent 95% confidence interval. **E.** Growth of wild type, Δ*hupA,* Δ*hupS* and Δ*hupA*Δ*hupS s*trains on solid SFM medium after exposure of spores to high temperature (60°C) or detergent (5% SDS) for 30 minutes before plating. **F.** Percentage of germinating spores of wild type (purple), Δ*hupA* (red)*, ΔhupS* (green) and Δ*hupAΔhupS* (blue) *s*trains after exposure to high temperature (60°C), detergent (5% SDS) or UV light.

Given the lack of spore pigmentation and chromosome segregation defects in the Δ*hupA*Δ*hupS* strain, we expected that its spores would be less viable than those of Δ*hupA* or Δ*hupS* spores. Indeed, spores from the Δ*hupA*Δ*hupS* strain were significantly less resistant to all tested stress factors: high temperature, presence of SDS as well as exposure to UV light or hydrogen dioxide. Colony forming unit (CFU) calculations showed that less than 0.00001% of Δ*hupA*Δ*hupS* spores survived a 45 minutes incubation in 60°C compared to survival of 40.5% of wild type spores and 1.3% and 0.07% of Δ*hupA* and Δ*hupS* spores, respectively. Treatment with SDS affected solely the spores of Δ*hupA*Δ*hupS* strain (less than 1% spores germinating), while for wild type and single deletion strains, treatment with SDS increased the germination rate from 60% to around 100% (Fig. 1 E, F). UV light exposure for 60 seconds resulted in the survival of less than 0.1% of Δ *hupS* and Δ*hupA*Δ*hupS* spores, whereas spores of *ΔhupA* and wild type strains were less affected and 0.6 and 2% spores still formed colonies, respectively (Fig. 1 F). The presence of H_2_O_2_ in both liquid and solid medium strongly inhibited the growth of Δ*hupS* and Δ*hupA*Δ*hupS* strains, while Δ*hupA* strain tolerated higher concentrations of H_2_O_2._ The H_2_O_2_ concentration of 1.75 mM led to growth inhibition on Δ*hupA* strain but not wild type strain (Fig. S2). Thus, factors causing DNA damage such as UV light or hydrogen dioxide, were found to be particularly harmful to spores of strains lacking HupS.

In summary, the deletion of both genes encoding HU homologs, *hupA* and *hupS,* in *S. coelicolor* resulted in a more severe phenotype than that of the single deletion mutants: more pronounced growth retardation, chromosome segregation defects and changes in colony pigmentation. Severity of Δ*hupA*Δ*hupS* strain phenotype compared to single deletion mutants suggested a synergy between HupA and HupS. The absence of both HU homologs decreased also spores resistance to stress conditions. The elevated sensitivity to some of the stress factors, such as UV light or reactive oxygen species, could be explained either by lack of the physical protection of DNA or by transcriptional changes of genes involved in stress response.

### HupA and HupS regulons partially overlap

Given that diminished stress response may result from the impact of HU homologs on gene expression, we set out to test whether the elimination of *hupA* or *hupS* would lead to transcriptional changes in *S. coelicolor*. Since the lack of HupA and HupS in *S. coelicolor* resulted in somewhat different phenotypes, we expected that their regulatory networks may not entirely overlap. To establish the HupA and HupS regulons and compare them to transcriptional changes in the double mutant strain, we performed an RNA-seq experiment for the Δ*hupA*, Δ*hupS* and Δ*hupA*Δ*hupS* strains, in comparison to wild type control, all grown in liquid YEME/TSB medium. For each strain two timepoints were chosen based on the growth curves (Fig. S3): the middle of exponential growth (20 h for wild type strain, Δ*hupA* and Δ*hupS*, 24h for Δ*hupA*Δ*hupS* strain) and the early stationary phase of growth (26 h for wild type strain, Δ*hupA* and Δ*hupS*, 30h for Δ*hupA*Δ*hupS* strain). Differential expression analysis of all *S. coelicolor* genes using edgeR package determined which genes were affected by either *hupA* or *hupS* deletions when compared to wild type strain at the two tested time points (Table S3).

The number of genes whose transcription changed in D*hupA* was similar during exponential growth and early stationary phase (272 and 299 genes, respectively), while *hupS* deletion affected expression of more genes during exponential growth than in the stationary phase (431 and 140, respectively) (Fig. 2 A, B). Gene expression was most altered in the double deletion mutant, with 451 genes changed during exponential growth and 343 genes during the stationary phase (Fig. 2 A, B). Interestingly, in Δ*hupA*Δ*hupS* strain 213 genes were affected at both analysed time points (while in the D*hupA* and Δ*hupS* it was only 43 and 64 genes, respectively) (Fig. S4). This may suggest that double deletion strain did not undergo a distinct transition between growth phases or that it was still at an earlier stage of growth than either Δ*hupA* or Δ*hupS* strains.

**Figure 2.**
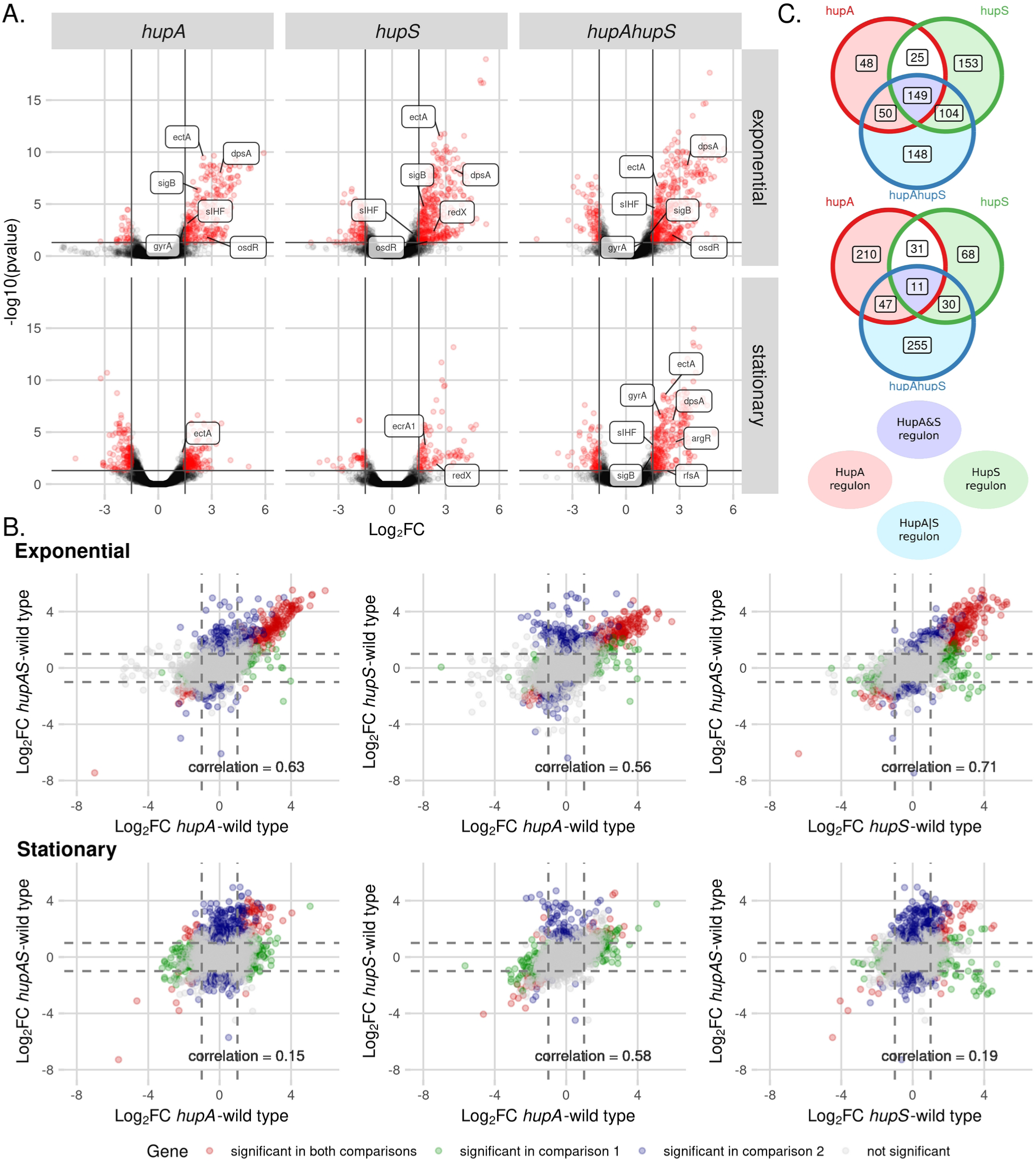
Global changes of gene expression in Δ*hupA*, Δ*hupS* and Δ*hupA*Δ*hupS* strains. **A.** Volcano plots showing altered gene expression in Δ*hupA, ΔhupS* and Δ*hupA*Δ*hupS* strains at exponential and stationary growth phase compared to the wild type strain. For each strain significantly changes genes (FDR ≤ 0.05, |Log_2_FC| > 1.5) are shown in red. Chosen differentially expressed genes are labelled. **B.** Venn diagrams showing significantly changed genes unique and common for analyzed strains; Δ*hupA, ΔhupS* and Δ*hupA*Δ*hupS* during exponential and stationary growth. **C.** Scatterplots showing correlation between Δ*hupA,* Δ*hupS* and Δ*hupA*Δ*hupS* transcriptomes from exponential and stationary growth compared to wild type strain. Pearson correlation coefficient is shown on each plot. Genes found to be significant in both comparisons (FDR ≤ 0.05, |Log_2_FC| > 1.5) are shown in red, while genes significant in only one of the comparisons are marked as blue or green, grey dots represent genes not affected by *hupA* and/or *hupS* deletions.

Most often, the deletion of *hupA* and/or *hupS* resulted in transcription upregulation (88% and 55% of genes in the Δ*hupA* mutant, 80% and 71% in the Δ*hupS* and 83% and 76% in the Δ*hupA*Δ*hupS* strains during exponential and stationary growth, respectively) (Fig. 2 A). To confirm the obtained results, ten representative genes from the putative *hupAS* regulon were chosen for replication analysis and their expression pattern was confirmed by an RT-PCR experiment (Pearson correlation coefficients 0.57 and 0.53 between Log_2_FC values for exponential and stationary growth, respectively) (Fig S5C).

To further compare the transcriptional changes between *hupA* and/or *hupS* deletion strains, we calculated the Pearson correlation coefficient of obtained Log_2_FC values from comparisons of mutant strains to the wild type strain (Fig. 2 B). We found a moderate correlation between the *hupA* and *hupS* strains at both stages of growth (Pearson coefficient = 0.56 and 0.58 at exponential and stationary growth, respectively). Interestingly, the transcriptional changes detected in the double deletion mutant Δ*hupA*Δ*hupS* were similar to those in the single deletion strains only during exponential growth (Pearson coefficient: 0.63 and 0.71 when compared to *hupA* and *hupS* strains, respectively), while during the stationary phase this strain was visibly distinct (Pearson coefficient: 0.15 and 019 when compared to *hupA* and *hupS* strains, respectively). The high similarity between RNA-seq results obtained for all deletion mutants at an earlier stage of growth and the remarkable difference of the Δ*hupA*Δ*hupS* strain from the other two strains in stationary phase could also be seen on the Principal Component Analysis (PCA) plot (Fig. S4, S5A).

Based on the pattern of regulation by HupA/HupS, differentially expressed genes could be divided into four major categories. The first group (HupA|S regulon) contained genes for which the presence of one HU homolog was sufficient to maintain the expression pattern. These genes were thus differentially expressed only in the Δ*hupA*Δ*hupS* strain – 148 and 255 genes during exponential and stationary growth, respectively. The second group (HupA&S regulon) required both HupA and HupS to maintain wild type level of expression. Therefore, this group was comprised of genes whose expression changed in all tested strains, 149 and 11 genes during exponential and stationary growth, respectively. The last two groups (HupA regulon and HupS regulon) contained genes whose expression changed in the Δ*hupS* strain, but not in the Δ*hupA* strain or in the Δ*hupA* strain, but not in the Δ*hupS* strain. Surprisngly, the HupS regulon was larger than the HupA regulon during exponential growth (257 and 98 genes, respectively) while HupA regulon was larger during stationary phase (Fig. 2C).

The larger number of genes under the control of HupA in the stationary phase and under control of HupS during exponential growth is somewhat contradictory to expectations based on the fact that HupA is the most abundant NAP during vegetative growth, while HupS levels increase during sporulation. The moderate overlap between HupA and HupS regulons and the existence of HupA|S and HupA&S regulons, indicates some extent of cooperation between these two HU homologs. This cooperation may explain their synergistic impact on phenotype, although the details of such cooperation remain to be elucidated. On the other hand, the separate HupA and HupS regulons may correspond to the different binding pattern of both proteins. While HupA was shown to predominantly bind within the central region of the *S. coelicolor* chromosome, HupS in *S. venezuealae* preferentially bound within the arms regions (29, 32). Finally, it is worth highlighting that both HU homologs seemingly most often played the role of transcriptional repressors. A similar function has been noted for Lsr2 in *S. venezeulae*, where removal of Lsr2 activated a number of secondary metabolite clusters (52). In *S. coelicolor,* a novel NAP called Gbn was also found to have a supressive effect on gene expression (53). The comparable roles of HupA and HupS positions them among other proteins crucial for controlling *Streptomyces* metabolism.

### Deletion of *hupA* and *hupS* genes alters the expression of genes involved in chromosome structure and topology maintenance

The fragility of Δ*hupA*Δ*hupS* strain spores, and to a lesser extent of the Δ*hupA* and Δ*hupS* strains, could be attributed either to lack of DNA protection by HU homologs or to transcriptional changes of genes vital for *S. coelicolor* spore maturation. Diminished resistance to stress conditions such as UV light, heat or free radicals has often been described for HU mutant strains of various bacterial species (33, 54–57). Therefore we investigated how the transcriptional activity of genes important for DNA structure and topology or spore maturation was changed in *hupA* and/or *hupS* deletion mutants (Fig. 3A).

**Figure 3.**
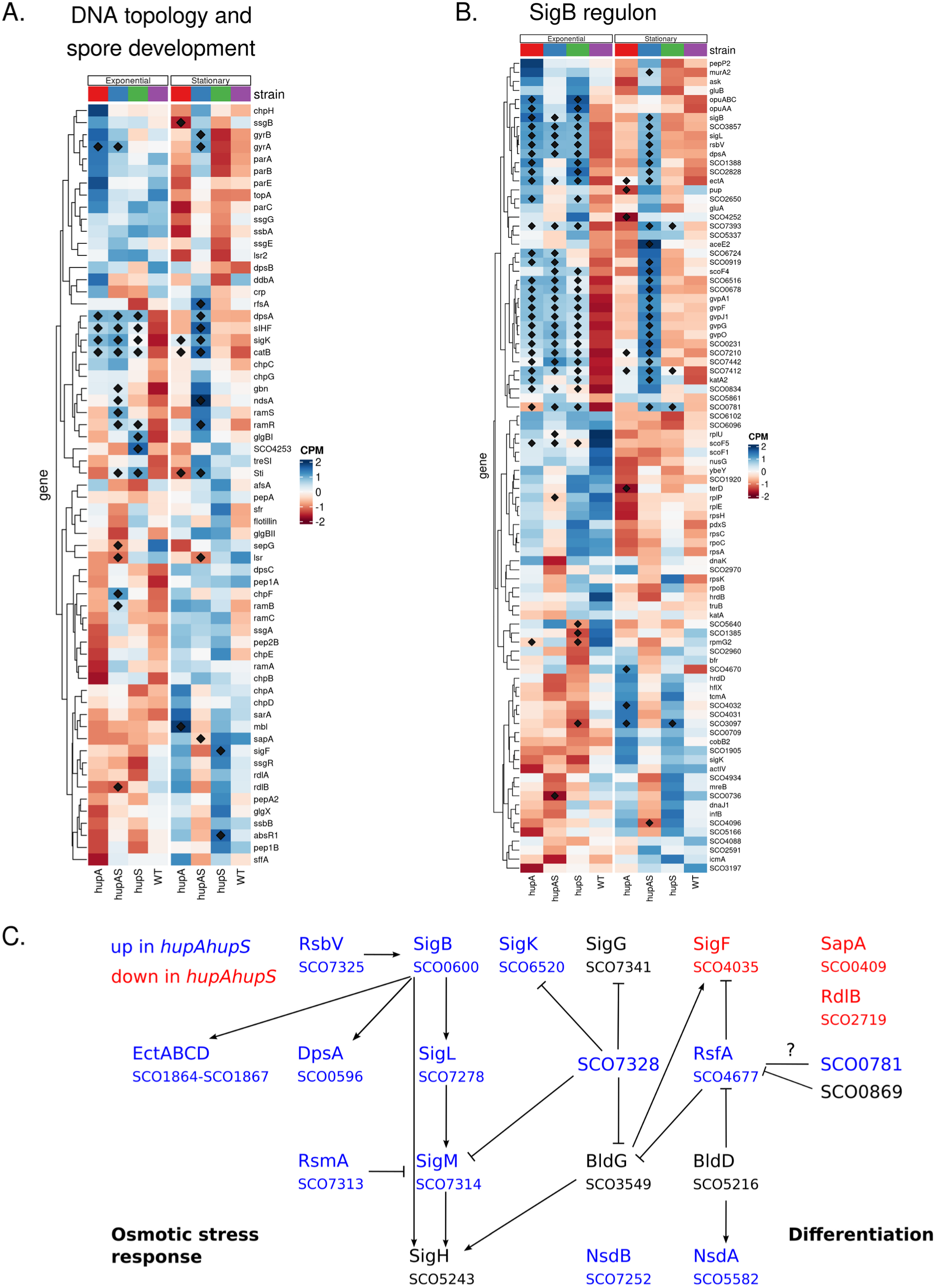
Deletion of *hupA* and/or *hupS* changes expression of genes connected to DNA topology maintenance and encoding global regulators. **A.** Heatmap showing normalized expression (CPM) of genes connected with DNA topology and spore development in wild type, Δ*hupA,* Δ*hupS* and Δ*hupA*Δ*hupS s*trains during exponential and stationary growth. **B.** Heatmap showing normalized expression (CPM) of SigB regulon genes in wild type, Δ*hupA,* Δ*hupS* and Δ*hupA*Δ*hupS s*trains during exponential and stationary growth. **C.** Selected genes from differentiation and stress response network that are upregulated (blue) or downregulated (red) in Δ*hupA*Δ*hupS* strain during stationary growth. **A, B** Genes marked with diamond were found to be significant in comparison to the wild type strain (FDR ≤ 0.05).

Since HupA is one of the most abundant NAPs in *S. coelicolor* during vegetative growth (30, 35), its deletion should be compensated by an upregulation of gene(s) encoding other NAP(s). Studies of *S. lividans* suggested that increased expression of *hupS* could partially suppress the effects of *hupA* deletion (36). However, we have not found any evidence of *hupS* upregulation in the Δ*hupA* strain or *hupA* upregulation in the Δ*hupS* strain. Instead, in both Δ*hupS* and Δ*hupA* mutants, we observed a significant increase in *dpsA* expression (*sco0596*), which encodes a NAP involved mainly in DNA protection in stressful conditions (58) and *sIHF* (*sco1480*), a NAP responsible for chromosome condensation and segregation (59, 60). Expression of *dpsA* and *sIHF* increased during exponential growth in all mutant strains as compared to the wild type (Log_2_FC for *dpsA* in Δ*hupA:* 3.45, in Δ*hupS:* 3.40 and in Δ*hupA*Δ*hupS:* 3.41; for *sIHF* in Δ*hupA:* 1.55, in Δ*hupS:* 1.24, in Δ*hupA*Δ*hupS:* 1.70), while in stationary phase expression of those genes increased only in the Δ*hupA*Δ*hupS* strain (Log_2_FC for *dpsA:* 2.61; for *sIHF*: 1.56) (Fig. 3 A, C). Deletion of *sIHF* was earlier shown to result in a phenotype similar to that of *hupA* and/or *hupS* mutants, namely reduced viability of spores and inhibition of sporulation (61). This suggests an existence of a functional overlap between HupA, HupS and sIHF in *S. coelicolor.* Interestingly, transcription of gene encoding another NAP named Gbn (*sco1839*) also increased in the Δ*hupA*Δ*hupS* strain (Log_2_FC Δ*hupA*Δ*hupS:* 1.75 and 1.31 during exponential and stationary growth, respectively) (Fig. 3 A). Deletion of *gbn* had a different effect then *hupAhupS* deletion and led to accelerated development, while overexpression of *gbn* delayed sporulation (53). Finally, unlike the above described NAP genes, the expression of *lsrL* (*sco4076*) decreased in the Δ*hupA*Δ*hupS* strain (Log_2_FC Δ*hupA*Δ*hupS* exponential: -1.24, stationary: -1.48). LsrL is a homolog of Lsr2, but little is known about its function in *Streptomyces*. Interestingly, the expression of *lsr2* remained unchanged in all tested strains.

HupA protein is also crucial for DNA supercoiling homeostasis in *S. coelicolor* and cooperates with topoisomerase I (TopA) in maintaining chromosome topology (32). Here, we did not detect any changes in expression of either *topA* (*sco3543*) or *parE/C* (*sco5822, sco5836*) genes encoding topoisomerase IV, but in strains with *hupA* deletion (*ΔhupA* and *ΔhupAΔhupS*), we observed an upregulation of the *gyrA/gyrB* operon (*sco3873-sco3874*) encoding gyrase, placing *gyrAB* in the HupA regulon (Fig. 3 A). This result corroborates an earlier observation that the D*hupA* mutant was more sensitive to gyrase inhibition by novobiocin than the wild type strain (32). Thus, the increased gyrase activity could be necessary to compensate for the lack of HupA.

Diminished UV and oxidative stress resistance of *hupA* and *hupS* spores could be linked to the role of HU in homologous replication or RecA-dependant DNA repair (56, 62). However, *recA* expression in *S. coelicolor hupA* and/or *hupS* mutants was not affected. Nevertheless, we found that some spore associated genes which were repressed in the Δ*hupA*Δ*hupS* strain during stationary growth, e.g., *sapA* (*sco0409*) and *rdlB* (*sco2719*) encoding spore coat proteins important for spore hydrophobicity (63–65) (Fig. 3 A, C). These changes could account for increased spores sensitivity to stress factors. Interestingly, even though mutant strain colonies were paler than the wild type, the genes responsible for spore pigmentation (e.g. *whiE*) were not affected.

Summarizing, genes encoding numerous proteins involved in DNA structure maintenance and protection, such as DpsA, sIHF and gyrase, were affected by either *hupA* or *hupS* deletion. This indicates at least partially independent functions of HupA and HupS in maintaining chromosome structure. Remarkably, in the exponential phase, the elimination of either HupA or HupS led to upregulation of expression, thus placing those genes in the HupA&S regulon. This suggests that in this phase of growth, in the absence of HupA and HupS, other NAPs may compensate for their loss and maintain chromosome organisation. However, in the stationary phase, the elimination of both HU homologs was required to activate other NAP encoding genes. This observation may be explained by modified chromosome structure during stationary phase (66). This may also explain the remarkably increased sensitivity of Δ*hupA*Δ*hupS* spores.

### HupA and HupS are a part of the *Streptomyces* regulatory network

The observed HupA- and HupS-dependent changes in global transcriptional activity might be explained by a direct impact of these NAPs on particular gene expression or by an indirect effect mediated by modified levels of regulators and/or sigma factors. To explore the latter possibility, we utilized the RNA-seq dataset to identify transcription regulators whose expression was altered in *hupA/hupS* deletion strains and whose regulatory networks have already been established. We found that genes encoding SigB (*sco0600*), ArgR (s*co1576*) and OsdR (*sco0204*) fulfilled these criteria. Next, we analysed how HupA- and/or HupS-dependent modification of these genes impacted their regulatory networks. We also investigated whether HupA and HupS are involved in the regulatory network controlling *S. coelicolor* life cycle.

SigB (*sco0600*) is a sigma factor, which acts as a major osmotic stress regulator and, through SigM (*sco7314*) and SigL (*sco7278*), regulates *Streptomyces* differentiation and stress response (*67*) (Fig. 3 B). The expression of *sigB* increased in Δ*hupA* and Δ*hupS* strains during exponential growth and in the Δ*hupA*Δ*hupS* strain during exponential and stationary growth (Fig. S5 B, C), placing *sigB* in the HupA&S regulon during exponential growth and in the HupA|S regulon during stationary growth. In *Streptomyces,* SigB activity is controlled by its anti-sigma factor RsbA (*sco0599*) and two anti-anti sigma factors: RsbB (*sco0598*) and RsbV (*sco7325*). Only *rsbV* expression was elevated in Δ*hupA* and/or Δ*hupS* strains. Out of 92 genes reported to belong to the SigB regulon about one-fourth were upregulated in the Δ*hupA*Δ*hupS* strain at both time points (25 genes, hypergeometric test p-value: 6.65*10^-12^) (Fig. 3 B). Remarkably, during exponential growth, most of these genes were also upregulated in either Δ*hupA* or Δ*hupS* deletion strains, while during stationary growth their expression was unchanged, reflecting the pattern of *sigB* expression levels. Genes from the SigB regulon that were upregulated by *hupA* and *hupS* deletions included *dpsA, sigL, sigM, ectABCD, rsbV* and *sco7590* (catalase) (Fig. 3 B, C). Thus, SigB network can serve as an example of HupA and HupS indirect influence, where changes of expression of a single sigma factor propagated through an entire regulatory network. However, a large fraction of genes from the SigB regulon were not found to be upregulated in neither *hupA* nor *hupS* deletion strains. This could be explained by either the low sensitivity of RNA-seq method to small changes in gene expression or by the influence of other factors independent of *hupA* and/or *hupS* deletion.

Another global regulator that was affected by the double deletion of *hupA* and *hupS* genes was ArgR. The expression of *argR* (*sco1576*) increased during stationary growth in Δ*hupA*Δ*hupS* strain (Log_2_FC Δ*hupAΔhupS*: 2.69), placing it in the hupA|S regulon. According to published data, ArgR controls the expression of around 1500 genes and usually acts as a repressor (68), but for our analysis, we only considered 90 genes whose expression was altered by *argR* deletion at all tested time points. Surprisingly, we found that the upregulation of *argR* in the Δ*hupA*Δ*hupS* strain was accompanied by upregulation of ∼ 30 genes belonging to ArgR cluster. These changes were observed during exponential growth of all analyzed mutant strains and in the Δ*hupA*Δ*hupS* strain during stationary growth (hypergeometric test p-value: 1.15*10^-16^). (Fig S6 A). In *E. coli,* HU proteins act as co-repressors with the GalR protein and *hupAB* deletion leads to the destabilization of repression loops and expression of the *gal* operon (69, 70). A similar mechanism could perhaps explain the observed upregulation of the *arg* operon despite the increased expression of the *argR* repressor gene found in the Δ*hupAΔhupS* strain.

In Δ*hupA, ΔhupS* and *ΔhupAΔhupS* mutants, we found a significant increase in the expression of genes encoding two-component system OsdKR (s*co0203-sco0204*) which thus fall into the HupA&S regulon (exponential growth, Log_2_FC Δ*hupA:* 1.44 and 2.65, *ΔhupS:* 1.28 and 2.07, Δ*hupAΔhupS:* 1.41 and 2.50). OsdR plays an important role in the control of stress and development related genes, and is an orthologue of *Mycobacterium tuberculosis* DevR protein (71, 72). The core regulon of OsdR/K system lies between genes *sco0167* and *sco0219.* These genes were shown to be activated by OsdR and involved in stress response, spore maturation and nitrogen metabolism in *S. coelicolor* (*71*). In all tested *hupA* and/or *hupS* strains, the expression of the “core” OsdR/K genes increased during exponential growth, but genes belonging to the OsdR regulon located in other parts of *S. coelicolor* chromosome were not affected (Fig. S6B). Interestingly, 17 genes belonging to the OsdR regulon were earlier shown to be influenced by TopA depletion (35), suggesting that this cluster could be controlled by DNA supercoiling. The increased expression of genes belonging to the *osdR* regulon during early stationary growth in all tested strains is in agreement with previously published transcriptomic data (31, 71).

Lastly, we observed an induction of *rfsA* gene (*sco4677*) but only in the Δ*hupA*Δ*hupS* strain (stationary growth, Log_2_FC: 2.27). This gene encodes an anti-sigma factor interacting with SigF. *rfsA* null mutants are characterized by faster development (73, 74). Additionally, RfsA is able to negatively regulate BldG (*sco3579*) by phosphorylation, linking it to the *Streptomyces* sporulation regulatory network (75). Interaction of RfsA with two anti-anti-sigma factors was described (74), but only one of them (*sco0781*) was induced in both strains lacking *hupS.* Expression of *sigF (sco4035)* decreased in the *ΔhupAΔhupS* strain during stationary growth. Though this result was not highly significant (FDR = 0.064), it could explain the spore fragility of *ΔhupAΔhupS* since *sigF* null mutants in *Streptomyces* are characterized by lessened resistance to detergent treatment (76, 77). Moreover, another anti-sigma factor, *sco7328,* was upregulated in the Δ*hupA*Δ*hupS s*train. This protein similarly to RfsA phosporylates BldG, but also inhibits activity of SigM, SigG (*sco7341*) and SigK (78) (Fig. 3 C). The expression pattern of *rsfA*, *sigF* and *sco7328* places them in the HupA|S regulon.

To sum up, the analysis of SigB regulon represents an example of a predictable secondary impact of *hupA* and *hupS* deletion resulting from *sigB* upregulation. In contrast, the ArgR regulon indicates the involvement of HU homologs in more complex regulatory circuits, while OsdR regulon expression could be influenced by structural changes of *S. coelicolor* chromosome caused by either *hupA* or *hupS* deletion. On the other hand, RsfA upregulation could partially explain the slower growth of the Δ*hupAΔhupS* strain.

### Deletion of *hupA* and/or *hupS* affects production of secondary metabolites

Given that the chromosome of *S. coelicolor*, similarly to other *Streptomyces* species, encodes numerous biosynthetic gene clusters, we set out to examine if the deletion of *hupA* and/or *hupS* in *S. coelicolor*, like *lsr2* deletion in *S. venezuelae,* could activate those clusters. We found that out of 22 secondary metabolic clusters present in *S. coelicolor* (Bentley, 2002) 4 exhibited changes of gene expression in *hupA* and/or *hupS* deletion strains. The most striking example was the *red* cluster (*sco5877-sco5898*) encoding the red oligopyrrole prodiginine antibiotic - undecyloprodigiosin (79). Undecyloprodigiosins were suggested to have antimalarial and anticancer properties (80), and in *S. coelicolor* they were implicated in controlled cell death during development (81). Expression of the *red* cluster is under the positive control of pathway specific regulators RedD and RedZ and is upregulated during stationary phase, especially during growth in liquid media (31, 82, 83). The expression of almost entire *red* cluster was elevated in the Δ*hupS* strain (but not in the D*hupA* or D*hupA*D*hupS* strains, placing it in the HupS regulon) at both tested time-points as compared to the wild type (Log_2_FC range for the *red* cluster, Δ*hupS* stationary growth: 1.95 – 4.23), except for the transcription regulator *redZ* (s*co5881*). The *f*act that the double deletion of *hupA* and *hupS* did not lead to activation of the *red* cluster suggests that HupA presence is required for its expression. Indeed, *redZ w*as downregulated in the Δ*hupA* strain during stationary growth (Fig. 4 A). Additionally, two-component systems: *ecrA1/A2* (*sco2517-sco2518*) and *ecrE1/E2* (*sco6421-sco6422*) which are involved in transcriptional control of the *red* cluster (84, 85) were also upregulated during stationary growth in Δ*hupS* strain. Notably, the overexpression of *red* cluster was earlier observed in the strain with deletion of *sIHF* (*60, 61*). Overproduction of RED antibiotic was confirmed by plate cultures showing an abundance of red pigment produced by the Δ*hupS* strain (Fig. 4 B). Interestingly during growth on solid medium, Δ*hupA* and Δ*hupA*Δ*hupS* strains did not produce the characteristic for *S. coelicolor* blue pigment (actinorhodin) (Fig. 4 B), which could be explained either by the growth delay or transcriptional influence of *hupA* deletion on the actinorhodin cluster transcription. However our RNA-seq data did not show any significant changes in the *act* cluster genes expression (Fig. S7). To sum up, HupA is required for the activation of gene encoding activator RedZ while HupS inhibits *red* cluster expression possibly by downregulation of the two component system genes.

**Figure 4.**
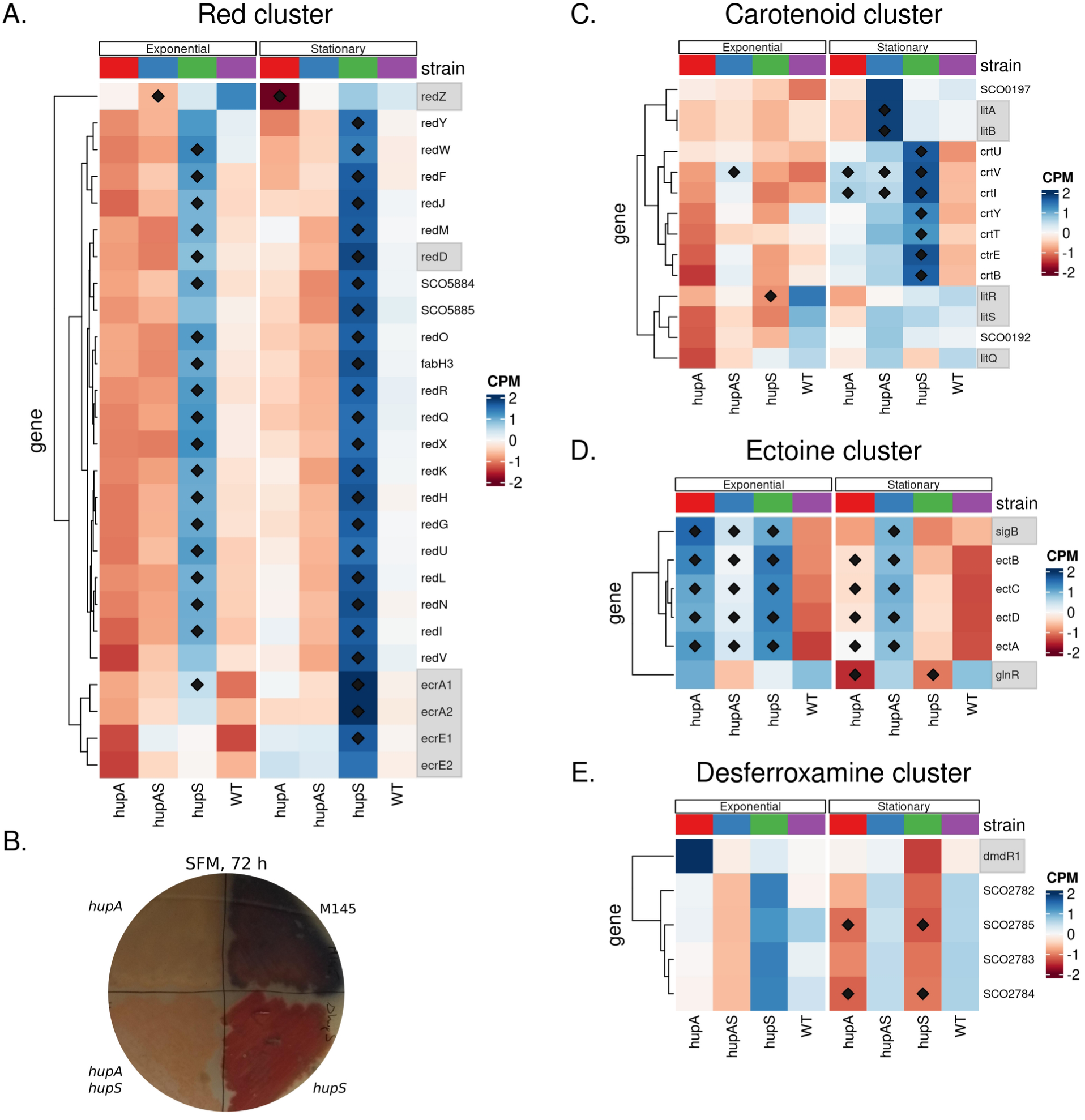
Deletion of *hupA* and/or *hupS* changes expression of genes associated with production of secondary metabolites. **A.** Heatmap showing normalized expression (CPM) of the red cluster genes in wild type, Δ *hupA,* Δ*hupS* and Δ*hupAΔhupS s*trains during exponential and stationary growth. **B.** Production of undecyloprodigiosin (red metabolite) during plate culture of wild type, Δ*hupA,* Δ*hupS* and Δ*hupA*Δ*hupS s*trains incubated of solid SFM medium for 72 hours in 30°C. **C.** Heatmap showing normalized expression (CPM) of the carotenoid gene cluster in wild type, Δ*hupA,* Δ*hupS* and Δ*hupA*Δ*hupS* strains during exponential and stationary growth. **D.** Heatmap showing normalized expression (CPM) of ectoine gene cluster in wild type, Δ*hupA,* Δ*hupS* and Δ*hupA*Δ*hupS* strains during exponential and stationary growth. **E.** Heatmap showing normalized expression (CPM) of desferroxamine cluster genes in wild type, Δ*hupA,* Δ*hupS* and Δ*hupA*Δ*hupS* strains during exponential and stationary growth. **A, C, D, E** Genes marked with diamond were found to be significant in comparison to the wild type strain (FDR ≤ 0.05). Genes encoding established transcription regulators are marked in grey.

The other biosynthetic gene cluster affected by the elimination of HupS was the carotenoid cluster (*sco0185-sco0194*). Expression of the *crt* cluster was shown to be highest during exponential growth (31). The function of carotenoids in *Streptomyces* has not been fully established yet, but it is suggested that they are involved in protection from photo-oxidative damage (86). Expression of the *crt* cluster was upregulated strongly in the Δ*hupS* strain during stationary growth (Log_2_FC range for *crt* cluster = 2.09 - 4.54) and to a lesser degree in the Δ*hupA* (Log_2_FC = -0.05 - 2.72) and Δ*hupA*Δ*hupS* (Log_2_FC = 0.42 - 2.24) strains (Fig. 4 C), placing it in the HupA&S regulon. Interestingly, genes belonging to the *crt* cluster are located between genes belonging to the *osdR* regulon, which expression was also elevated in either Δ*hupA* or Δ*hupS* mutants during exponential growth (Fig S6). Expression of the *crt* cluster is light induced and controlled by *litQR (sco0193-sco0194)* and *litSAB* genes, with *litS* being essential for *crt* expression *(sco0195-sco0197*) (Takano, 2005). Notably, the expression of *litSAB* genes in Δ*hupA* and/or Δ*hupS* strains was not significantly different from the wild type strain with the exception of *litA,* which was upregulated in Δ*hupA*Δ*hupS* strain during stationary growth (Fig. 4 C).

Elimination of HupA and/or HupS activated the ectoine cluster (*sco1864-sco1887)*. Ectoine production in *Streptomyces* has been linked with survival in high salt or temperature conditions (87). Earlier transcriptional studies showed that the *ect* cluster expression is highest during exponential growth and decreases at later stages of growth on both solid and liquid media (31, 82, 88). Deletion of *hupA* and/or *hupS* genes led to an overexpression of *ectABCD* genes at both analysed time points (Log_2_FC range for *ect* cluster exponential growth, *ΔhupA:* 2.54 - 2.94; *ΔhupS:* 2.68 - 4.00; *ΔhupAΔhupS:* 1.37 – 2.15) thus placing this cluster in the HupA&S regulon (Fig. 4 D). The ectoine cluster was shown to be negatively controlled by the GlnR transcription factor (*sco4159*) (89) and expression of *glnR* decreased in Δ*hupA* and Δ*hupS* strains during stationary growth. Additionaly *ect* cluster expression is dependent upon SigB (67, 90), and expression of *sigB* was elevated in all three *hupA* and/or *hupS* deletion strains (Fig. 4 D). Thus, ectoine cluster activation may be explained by increased levels od SigB and lowered levels of GlnR in the absence of HU homologs.

Contrary to the previously described clusters, the desferroxamine cluster *(sco2782-sco2785)* was downregulated in the absence of HupA or HupS. The *des* cluster encodes genes necessary for the production of desfeeroxamine, a nonpeptide hydroxamate siderophore (91). Desferrioxamine are produced by many *Streptomyces* species in iron deficiency conditions and at an early stage of growth on solid media (88, 92). Expression of *desABCD* genes decreased in Δ*hupA* and Δ*hupS* strains during stationary growth (Log_2_FC range Δ*hupA:* -2.24 to -3.08; Δ*hupS:* -2.65 to -3.30), but not in the double deletion Δ*hupA*Δ*hupS* strain (Fig. 4E). *desABCD* expression is governed by the iron repressor *dmdR1 (sco4394)* (*93*) but *dmdR1* transcription was not affected by *hupA* and/or *hupS* deletion (Fig. 4 E). Thus, the mechanism of *des* cluster regulation by HupA and HupS is unclear.

To sum up, we found that expression of four BGC was modified in the absence of HupA and/or HupS. Expression of secondary clusters responsible for the production of RED and carotenoid compounds were activated by *hupS* deletion, however the presence of HupA was required for this activation. That suggests an interplay between both HU homologs in the transcriptional regulation of these clusters. In contrast, ectoine cluster was upregulated in the absence of at least one HU homolog. For three out of four clusters, the changes of expression could be at least partially explained by modified levels of the pathway specific (RedZ) or global (SigB) regulators.

### Effective response to osmotic stress depends upon the presence of HupA

Several sigma factors whose expression levels were affected by *hupA* and/or *hupS* deletion participate in *S. coelicolor’s* response to osmotic stress (i.e. SigB). Similarly, ectoine, whose biosynthesis gene cluster expression was elevated in *hupA* and/or *hupS* deletion mutants, is important for *Streptomyces* survival in high salt environments (87). These observations prompted us to determine the effect of *hupA* and/or *hupS* deletion on *S. coelicolor* survival and gene expression in osmotic stress. To this end, we cultured Δ*hupA,* Δ*hupS* and Δ*hupA*Δ*hupS* strains in liquid YEME/TSB medium supplemented with 0.5 M NaCl, to assess their growth rate, and next, we to determine the transcription profile using RNA-seq (Table S3).

Addition of 0.5 M NaCl slowed down growth of all tested strains, but the inhibition of growth was most pronounced for Δ*hupA* and Δ*hupA*Δ*hupS* strains (Fig. S3). NaCl supplementation altered expression of a substantial number of genes in the wild type strain, with more genes affected during the stationary than the exponential phase (142 and of 338 genes, respectively). Deletion of *hupA* or *hupS* also resulted in a significant change in transcryptomic activity compared to these strains cultured in normal medium (659 and 455 genes, respectively during stationary growth). However, the double deletion mutant was largely unaffected by osmotic stress, with only 95 changed genes during stationary growth in the NaCl-supplemented medium (Fig. 5 A). Next, we calculated the correlations between individual strains’ responses to the osmotic stress. The Δ*hupS* strain transcriptome was the most similar to the wild type strain (Pearson coefficient = 0.71), while Δ*hupA* and *ΔhupAΔhupS* strains were significantly different from the wild type strain (Pearson coefficient 0.25 and 0.31, respectively). Both strains lacking the *hupA* gene were remarkably similar to each other with a 0.79 correlation coefficient and 432 shared genes (Fig 5B, C, S8).

**Figure 5.**
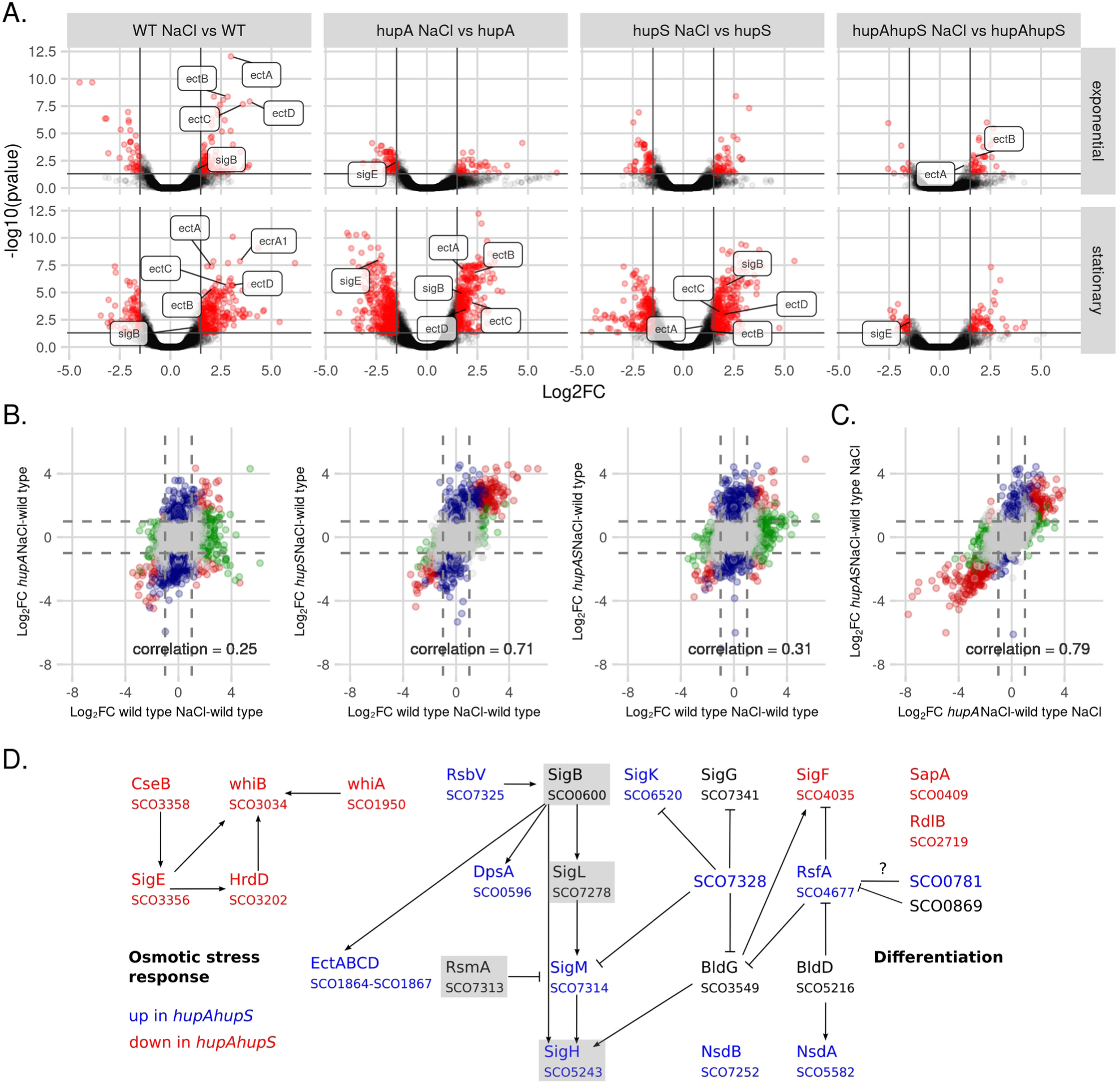
*hupA* deletion changes *S. coelicolor* response to growth in osmotic stress. **A.** Volcano plots showing changes in gene expression in wild type, Δ*hupA, ΔhupS* and Δ*hupAΔhupS* strains from exponential and stationary phase of growth in NaCl supplemented medium (0.5 M NaCl) compared to the same strain grown in standard YEME/TSB medium. For each strain significantly changes genes (FDR ≤ 0.05, |Log_2_FC| > 1.5) are shown in red. Chosen differentially expressed genes are labeled. **B.** Scatterplots showing correlation between wild type and Δ*hupA, ΔhupS* and Δ*hupAΔhupS* Log_2_FC values in medium supplemented with salt. Log_2_FC values were calculated from comparison with wild type strain grown in standard medium. Pearson correlation coefficient is shown on each plot. Genes found to be significant in both comparisons (FDR ≤ 0.05, |Log_2_FC| > 1.5) are shown in red, while genes significant in only one of the comparisons are marked as blue or green, grey dots represent genes not affected by *hupA* and/or *hupS* deletions. **C.** Correlation between Δ*hupA* and Δ*hupAΔhupS* transcriptomes during stationary growth in NaCl supplemented medium. Pearson correlation coefficient is shown on the plot. Genes found to be significant in both comparisons (FDR ≤ 0.05, |Log_2_FC| > 1.5) are shown in red, while genes significant in only one of the comparisons are marked as blue or green, grey dots represent genes not affected by *hupA* and double *hupAhupS* deletions. **D.** Regulatory networks network involving selected upregulated (blue) or downregulated (red) genes in Δ*hupA*Δ*hupS* strain during stationary growth in NaCl supplemented medium (0.5 M NaCl).

These observations suggest that the presence of HupA is necessary for *S. coelicolor* response to osmotic stress. This situation could partly explained by the fact that several genes overexpressed in the wild type strain in NaCl supplemented medium, like *ectABCD* or *sigB,* are a part of the HupA&S regulon (Fig. 5 A). Moreover, among the genes most affected by *hupA* deletion, we found *sigE (sco3356);* a sigma factor related to osmotic stress (94, 95). SigE controls the expression of genes mainly involved in maintaining cell wall and membrane integrity (95). HupA could be one of SigE binding partners (96). Lack of SigE could explain some of the transcriptomic changes observed in both Δ*hupA* strains. Indeed, analysis of the SigE regulon revealed the presence of 42 genes that were strongly repressed in either Δ*hupA* or *ΔhupAΔhupS* strains but induced in the wild type and Δ*hupS* strains during osmotic stress (hypergeomertic test p-value: 2.49 * 10^-14^) (Fig. S8 D). On the other hand, *in vitro* studies concerning the HupA protein showed that its binding to DNA is affected by NaCl concentration (10). Perhaps altered distribution of HupA on the *S. coelicolor* chromosome is one of the sources of observed transcriptional alterations.

### Gene clusters within HupAS regulons

Lastly, to improve gene classification into the four major regulons (HupA, HupS, HupA&S and HupA|S) by taking into account the influence of salt, we performed a cluster analysis of the entire dataset obtained for the four strains tested at two timepoints under two growth conditions. Using the *clust* programme we found 15 clusters of genes (with sizes ranging between 11 and 577) (Fig. S10), containing a total of 1601 *S. coelicolor* genes. Among the obtained clusters, three contained genes whose expression seemed to be affected by the lack of either of the HU homologs (clusters 9, 10 and 13) and thus belonging to the HupA&S regulon. Two of those clusters, 9 (39 genes) and 10 (142 genes), included many genes identified in the earlier differential expression analysis as controlled by both HupA and HupS, such as: *dpsA, sIHF, sigM, sigL, rsbV, catB, sigK* in cluster 10, and: *sigB, ectABCD* in cluster 9. The expression of genes from cluster 10 seemed to be largely unaffected by growth in osmotic stress, while genes belonging to cluster 9 were upregulated during stationary growth in osmotic stress conditions in all tested strains (Fig. 6). Interestingly, cluster 13 contained only genes located between 150 kb and 213 kb on the *S. coelicolor* genome and belonging to the OsdR regulon described earlier. Possibly, the control of cluster 13 expression involves changes in chromosome organization and/or supercoiling, explaining its sensitivity to both *hupA* and/or *hupS* deletion and TopA depletion (35) (Fig. 6).

**Figure 6.**
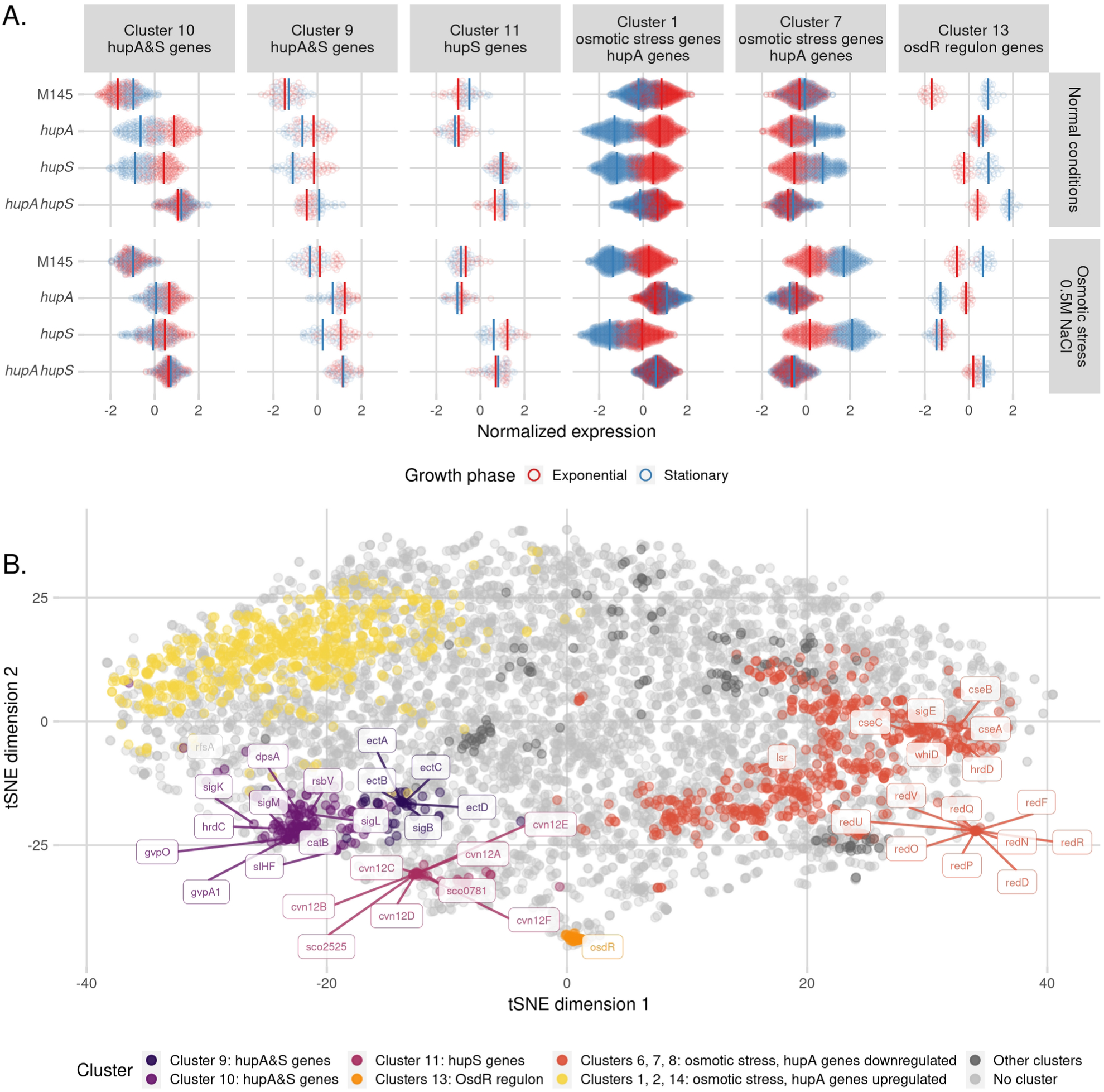
Clusters of genes within *hupAS* regulons. **A.** Cluster identification **-** results of clust analysis of wild type, Δ*hupA,* Δ*hupS* and Δ*hupA*Δ*hupS* transcriptomes from exponential (red) and stationary (blue) phase of growth in normal and NaCl supplemented medium. Plot shows six chosen clusters out of 15 obtained. **B.** t-distributed stochastic neighbour embedding (t-SNE) analysis of normalized wild type Δ*hupA,* Δ*hupS* and Δ*hupA*Δ*hupS* transcriptomes from exponential and stationary phase of growth in normal and NaCl supplemented medium. Clusters: HupA&S regulon (cluster 9 – dark purple, cluster 10 – purple, cluster 13 - orange), HupS regulon (cluster 11 – pink), HupA regulon clusters affected by growth in NaCl supplemented medium (clusters 6, 7, 8 – red, clusters 1, 2, 14 – yellow). Positions of genes associated with stress response, secondary metabolites production, spore development and NAPs are shown on the plot.

Cluster 11 (33 genes) contained genes whose expression changed only in Δ*hupS* or Δ*hupA*Δ*hupS* strains, thus constituting the HupS regulon. The expression of those genes was not influenced by growth in osmotic stress conditions and was also similar at both tested growth stages. Seemingly, those genes were controlled solely by HupS. This group included *cvn12* conservon (*sco2879-sco2884*), anti-anti sigma factor *sco0781,* putative LysR regulator (*sco2734*), putative stress response protein (*sco5806*), and operon *sco2521-sco2526.* SCO2525 is a putative methyltransferase necessary for normal growth of *S. coelicolor* (*97*) (Fig. 6). Genes affected only by *hupA* deletion (HupA regulon) belonged to six clusters (Clusters 1, 2, 6, 7, 8, 14), and their expression changed only during osmotic stress. Cluster 7 contained, among others, genes belonging to the *red* cluster (Fig. S9)*, sigE,* and *lsrL*. The expression of those genes increased during stationary growth in osmotic stress in the wild type and *ΔhupS* strain, but remained low in Δ*hupA* and *ΔhupAΔhupS* strains (Fig. 6, S10). This analysis confirmed that genes belonging HupA and HupA&S regulons are involved in osmotic stress response, while HupS regulon genes are not sensitive to increased salt concentration.

### Conclusions

In summary, the presence of both HupA and HupS is necessary for proper growth and development of *S. coelicolor*. The absence of both HupA and HupS results in severe growth inhibition and impaired stress survival, significantly more pronounced than that of either single deletions strains, indicating a synergy between the actions of the two HU homologs. The increased sensitivity of spores to DNA damaging factors may be explained by diminished protection of DNA.

RNA-seq analysis showed that by binding to DNA, HupA and HupS act as global transcription factors, altering the expression of multiple genes, mostly upregulating them. Genes upregulated in the absence of at least one of HU homologs were involved in DNA protection (NAPs, gyrase), transcription regulation (e.g. sigma factors) or stress survival (e.g. osmotic stress). HupS was involved in controlling the expression of secondary metabolites clusters (e.g. the *red* cluster), while HupA’s control of gene expression, separate from HupS, was mostly evident during growth in osmotic stress (Fig. 7). The identification of HupA&S and HupA|S regulons suggests a cooperation between the two HU homologs in *Streptomyces*.

**Figure 7.**
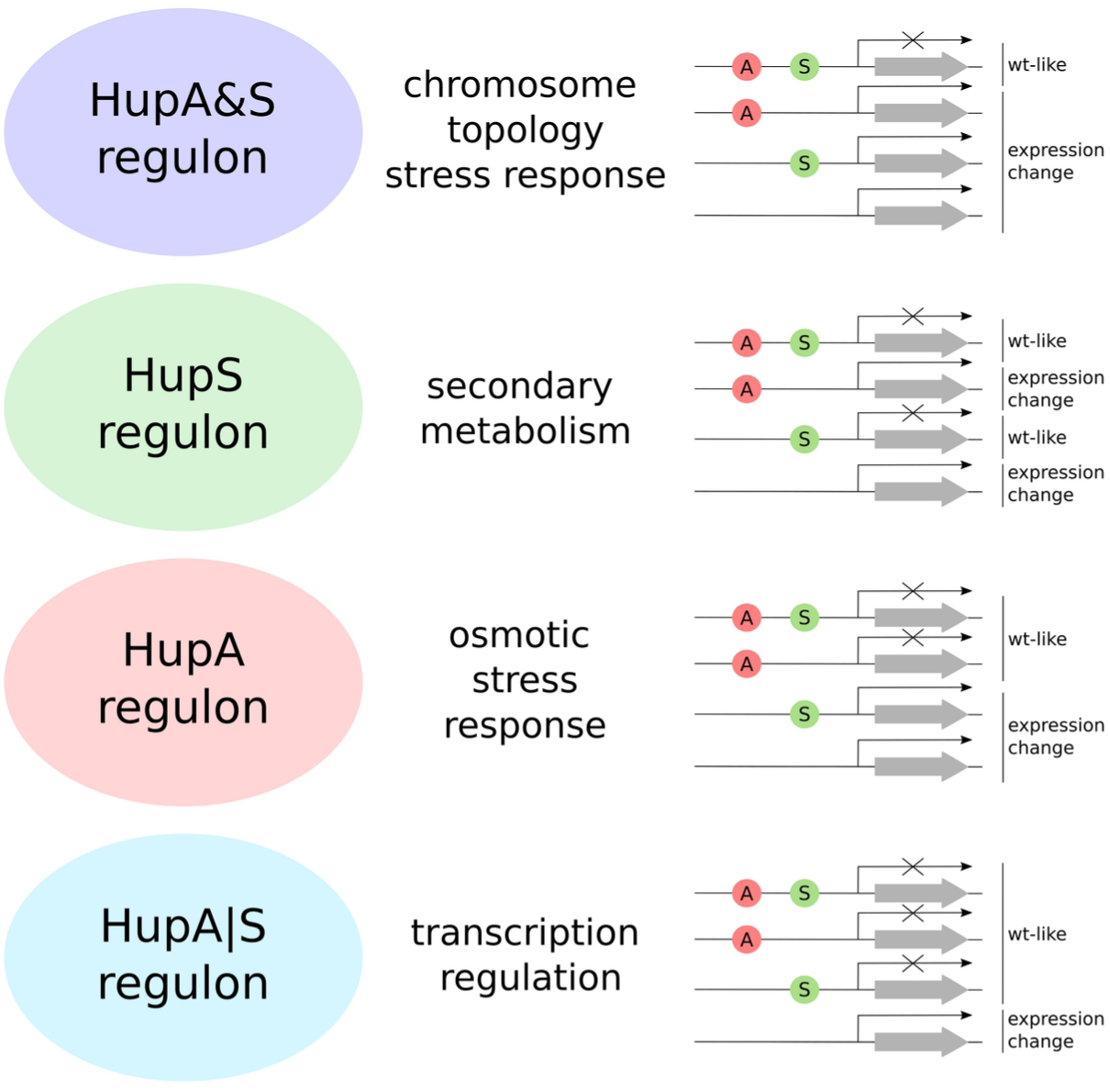
HupA and HupS regulons.

### Data Availability

The raw RNA-Seq data, as well as the processed data generated in this study, have been deposited in the ArrayExpress database (EMBL-EBI) under accession code E-MTAB-13846.

## Acknowledgements

We thank Klas Flärdh for the Δ*hupS* strain (K304). We thank Govind Chandra for help in bioinformatic analysis. We thank Mark Buttner for helpful collaboration.

## Funding

This work was funded by the Polish National Science Centre: SONATINA grant 2018/28/C/NZ1/00241.

## Supplementary Data

**Table S1.**
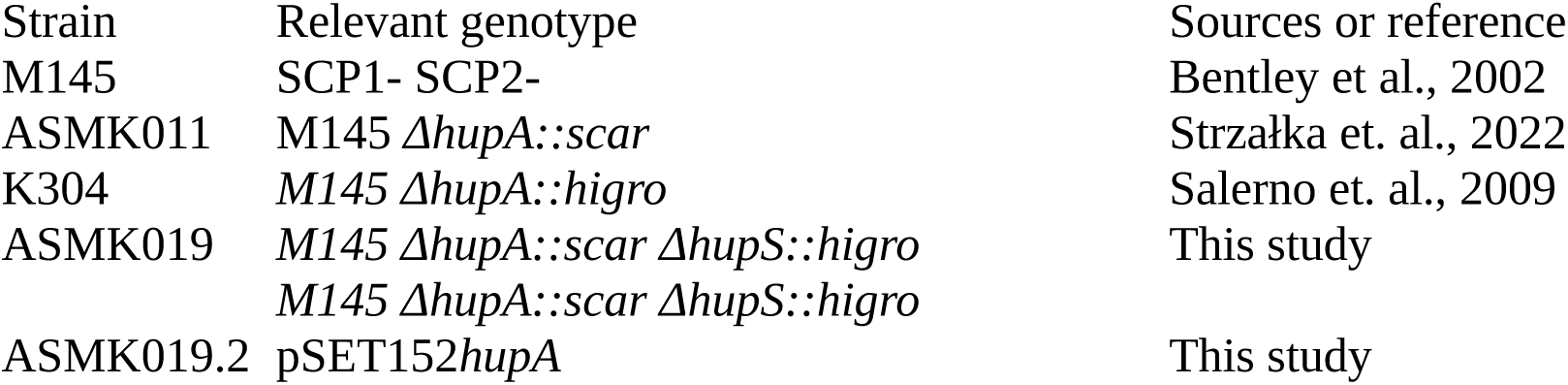
Streptomyces coelicolor strains used in the study.

**Table S2.**
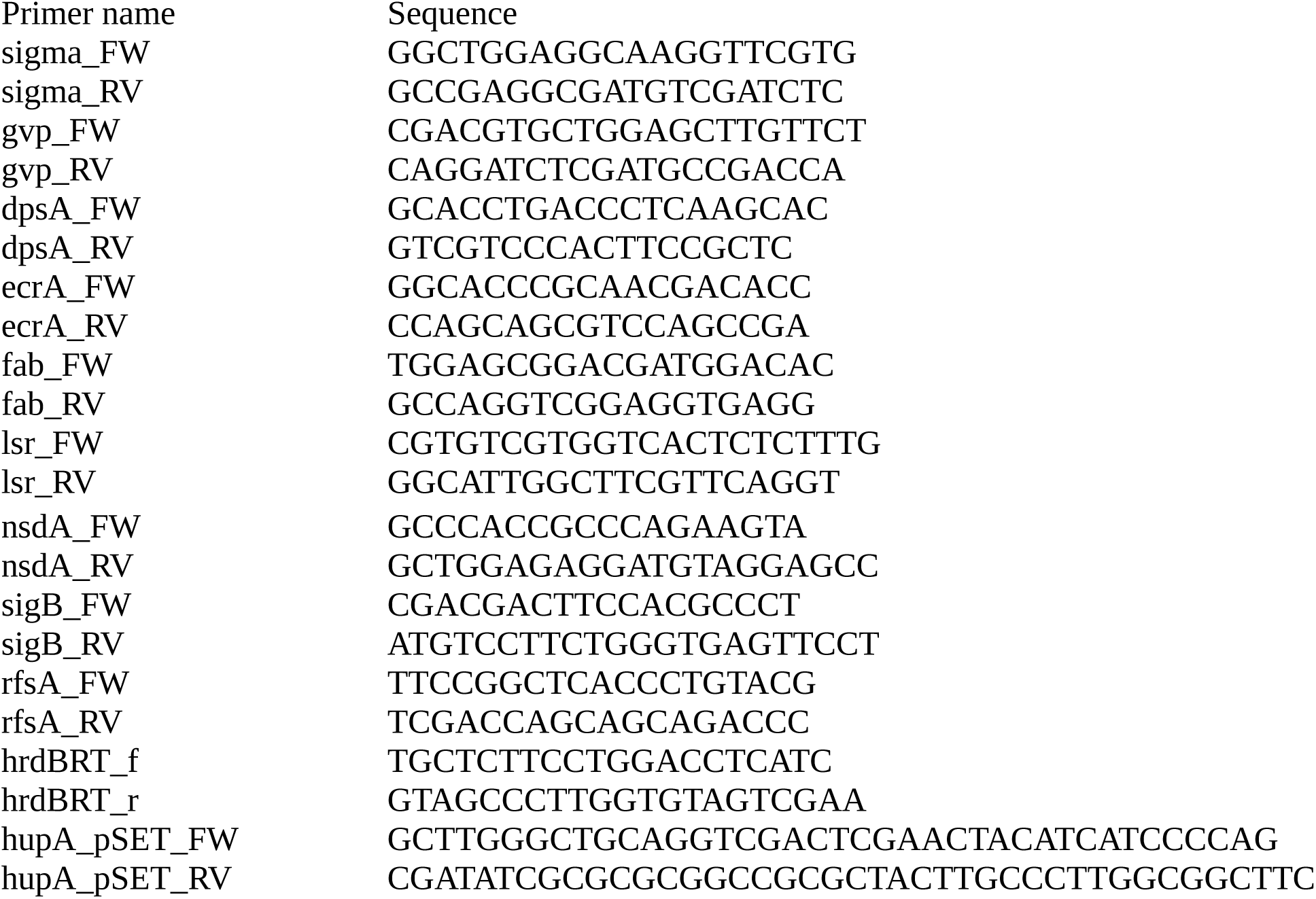
Oligonucleotides used in the study.

**Fig. S1.**
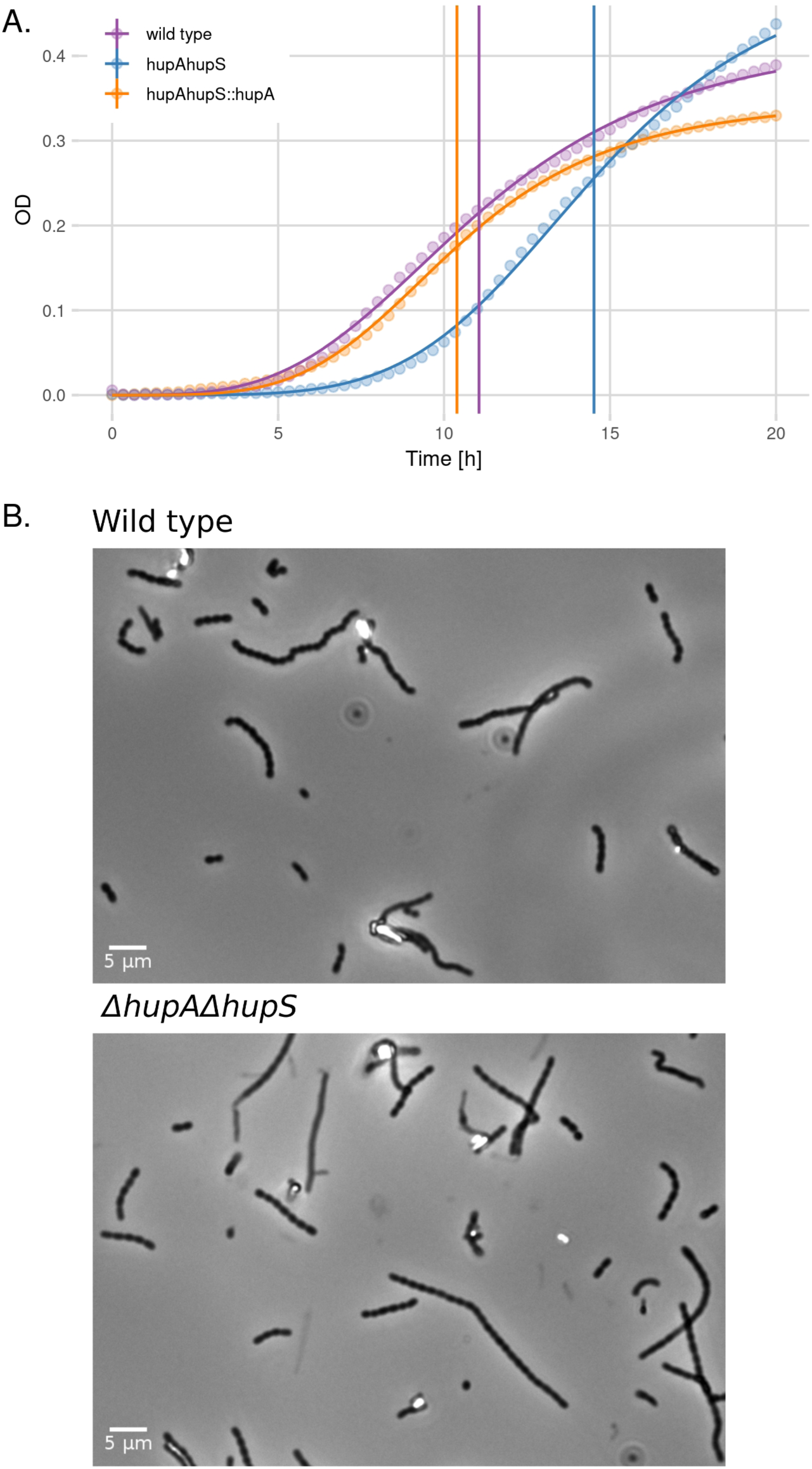
Growth and sporulation of Δ*hupA*Δ*hupS* strain. **A.** Growth curves of wild type (purple), Δ*hupA*Δ*hupS* (blue) and Δ*hupA*Δ*hupS::hupA* (orange) strains cultured in ‘79’ medium. Solid lines show the log-logistic model while vertical lines represent the half time calculated by the model. **B.** Microscopic images of spores of wild type *S. coelicolor* and Δ*hupA*Δ*hupS* strains from 7-day old colonies grown on SFM solid medium. Scale 5 μm.

**Fig. S2.**
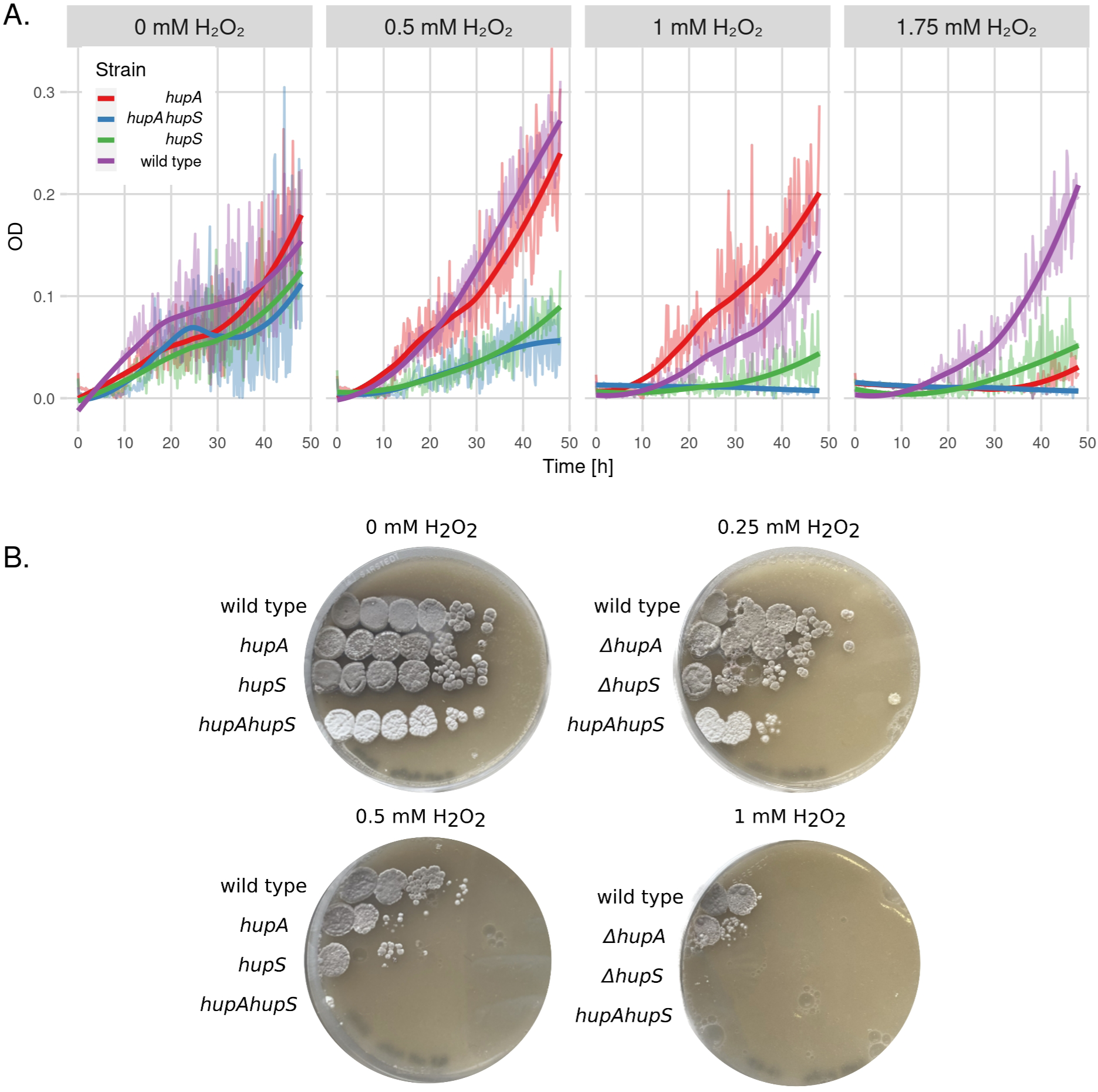
Impact of *hupA* and/or *hupS* deletion on *S. coelicolor* growth in the presence of H_2_O_2_. **A.** Growth curves of wild type (purple), Δ*hupA* (red), Δ*hupS* (green) and Δ*hupA*Δ*hupS* (blue) strains cultured in ‘79’ medium supplemented with increasing concentrations of H_2_O_2_. Solid lines show the growth curve smoothed by the *loess* algorithm. **B.** Growth of wild type, Δ*hupA,* Δ*hupS* and Δ*hupA*Δ*hupS* strains on solid SFM medium containing increasing concentrations of H_2_O_2_. Pictures were taken after 5 days of incubation in 30°C.

**Fig. S3.**
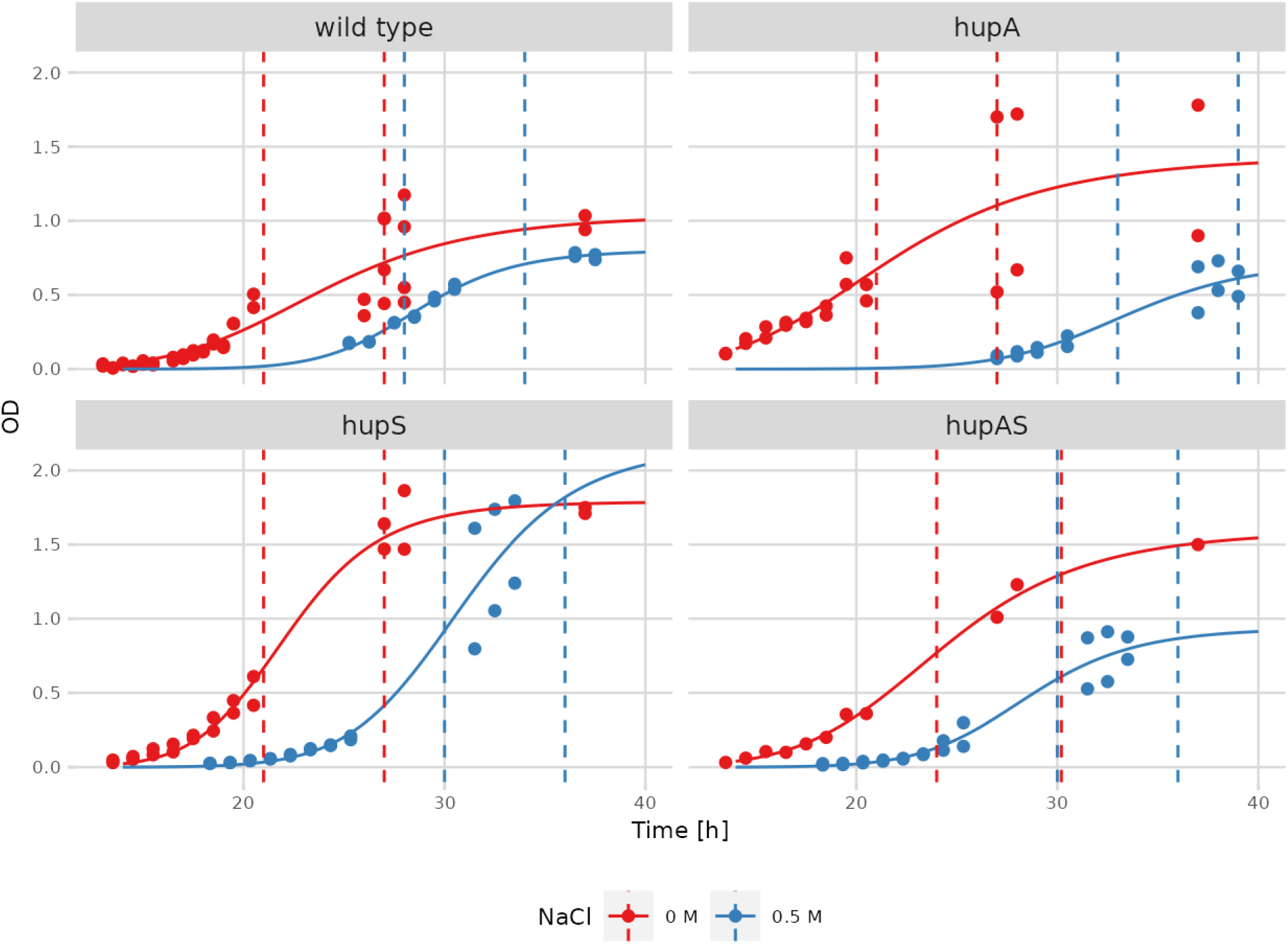
Impact of osmotic stress on the growth of wild type, Δ*hupA,* Δ*hupS* and Δ*hupA*Δ*hupS* strains. Growth curves of wild type, Δ*hupA,* Δ*hupS* and Δ*hupA*Δ*hupS* strains in standard (red) and NaCl supplemented (blue, 0.5 M NaCl) YEME/TSB medium. Vertical lines show growth half-time calculated by log-logistic model.

**Fig. S4.**
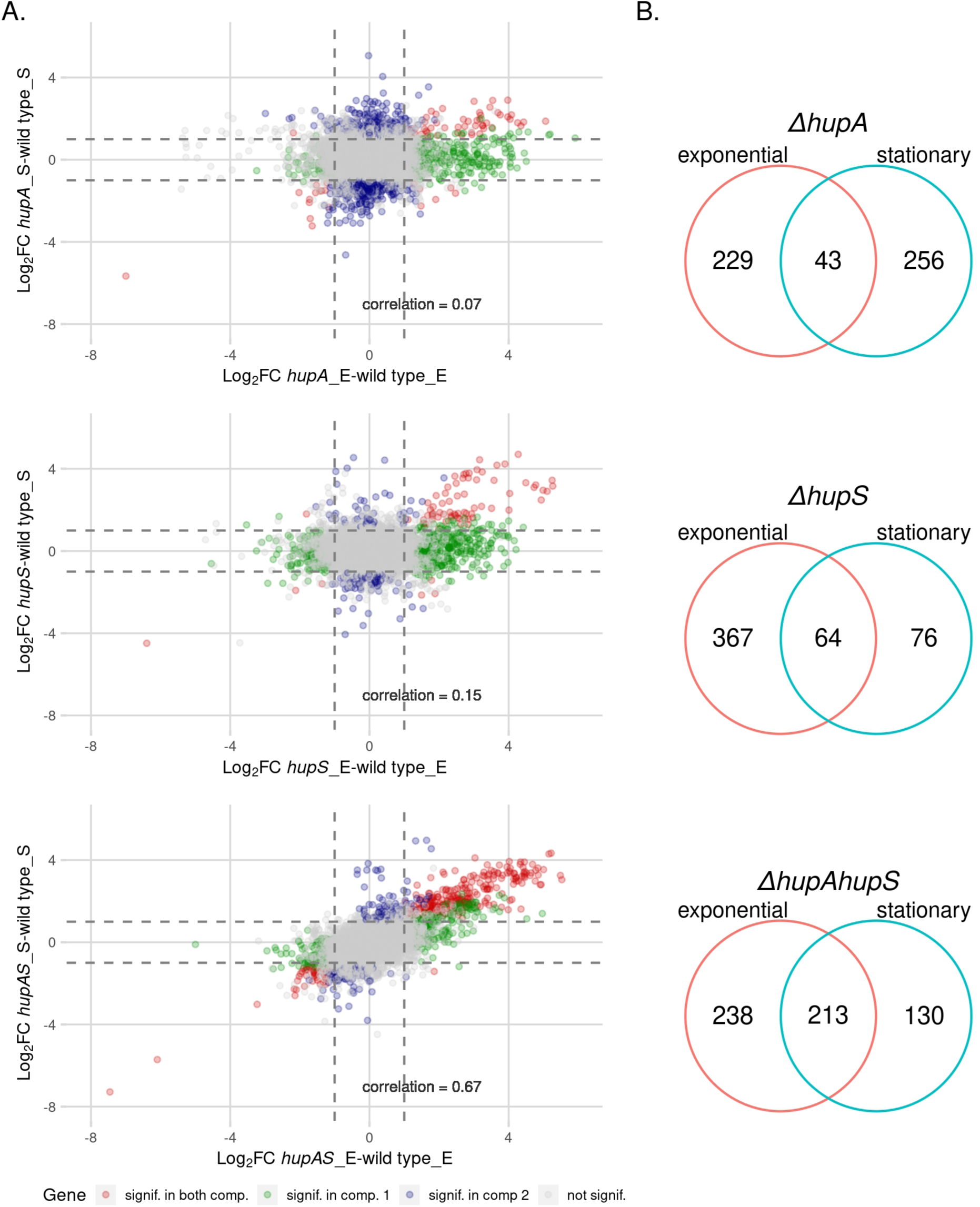
Comparison of transcriptomes of Δ*hupA,* Δ*hupS,* Δ*hupA*Δ*hupS* strains during different phases of growth. **A.** Scatterplots showing correlation between Δ*hupA,* Δ*hupS* and Δ*hupA*Δ*hupS* transcriptomes from exponential phase compared to stationary phase. Pearson correlation coefficient is shown on each plot. Genes found to be significant in both comparisons (FDR ≤ 0.05, |Log_2_FC| > 1.5) are shown in red, while genes significant in only one of the comparisons are marked as blue or green, grey dots represent genes not affected by *hupA* and/or *hupS* deletions. **B.** Venn diagrams showing significantly changed genes common for analyzed strains Δ*hupA, ΔhupS* and Δ*hupA*Δ*hupS* during exponential and stationary growth.

**Fig. S5.**
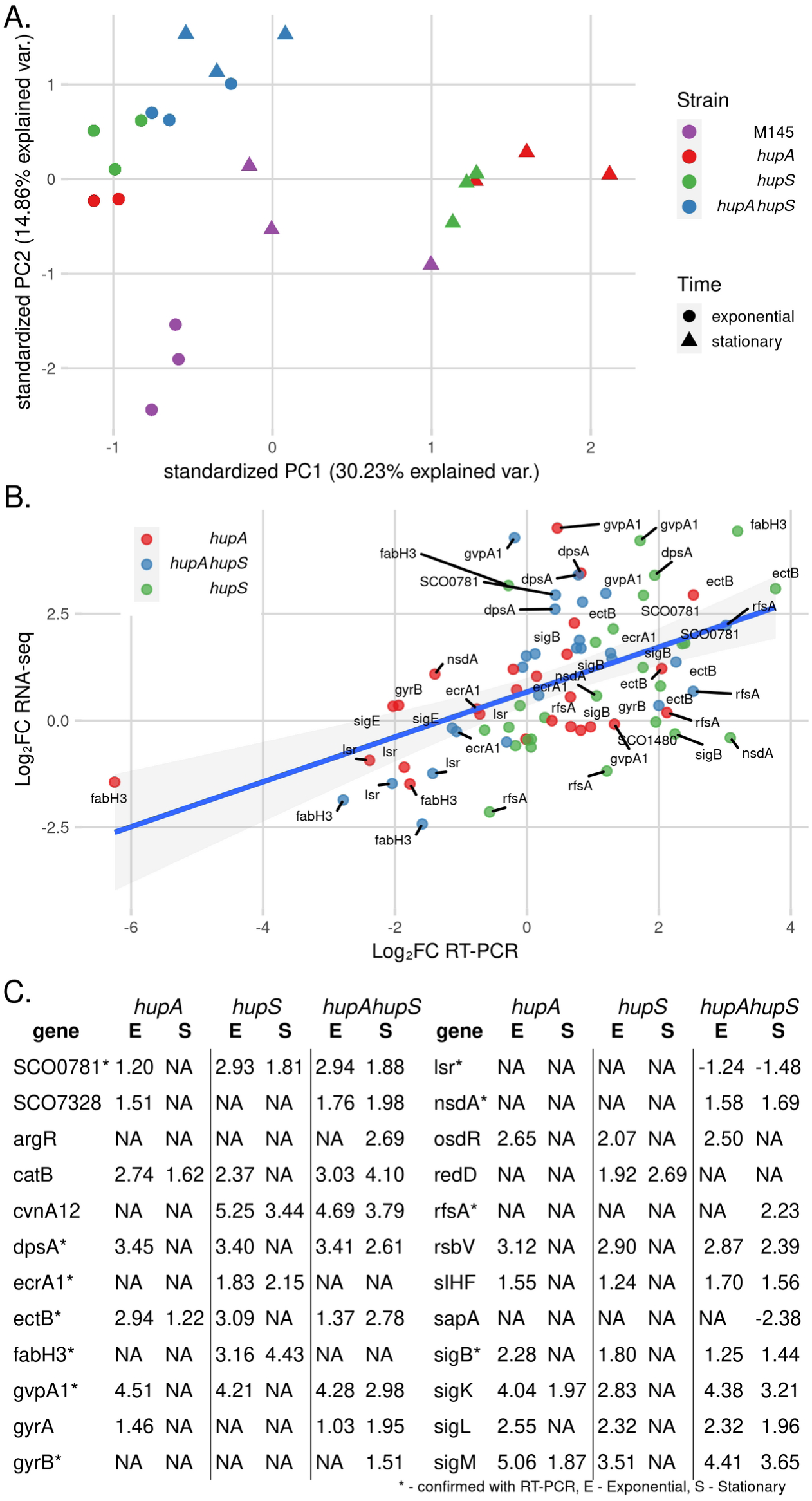
RNA-sea and RT-PCR analysis of gene expression. **A.** Principal component analysis (PCA) of the normalized RNA-seq CPM (Counts Per Million) data of *S. coelicolor* strains in response to *hupA* (red)*, hupS* (green) or *hupAhupS* (blue) deletion during exponential (circles) or stationary growth (triangles). **B.** Correlation between Log_2_FC values obtained from RNA-seq and RT-PCR for genes: *dpsA, sigB, gyrB, fabH3, ecrA1, sIHF, lsr, sco0781, gvpA1* from Δ*hupA* (red), Δ*hupS* (green) and Δ*hupA*Δ*hupS* (blue) strains when compared to the wild type strain. **C.** Table showing Log_2_FC values of several genes in strains Δ*hupA,* Δ*hupS* and Δ*hupA*Δ*hupS* when compared to wild type strain during exponential and stationary growth, NA value indicates that FDR value was above the 0.05 threshold.

**Fig. S6.**
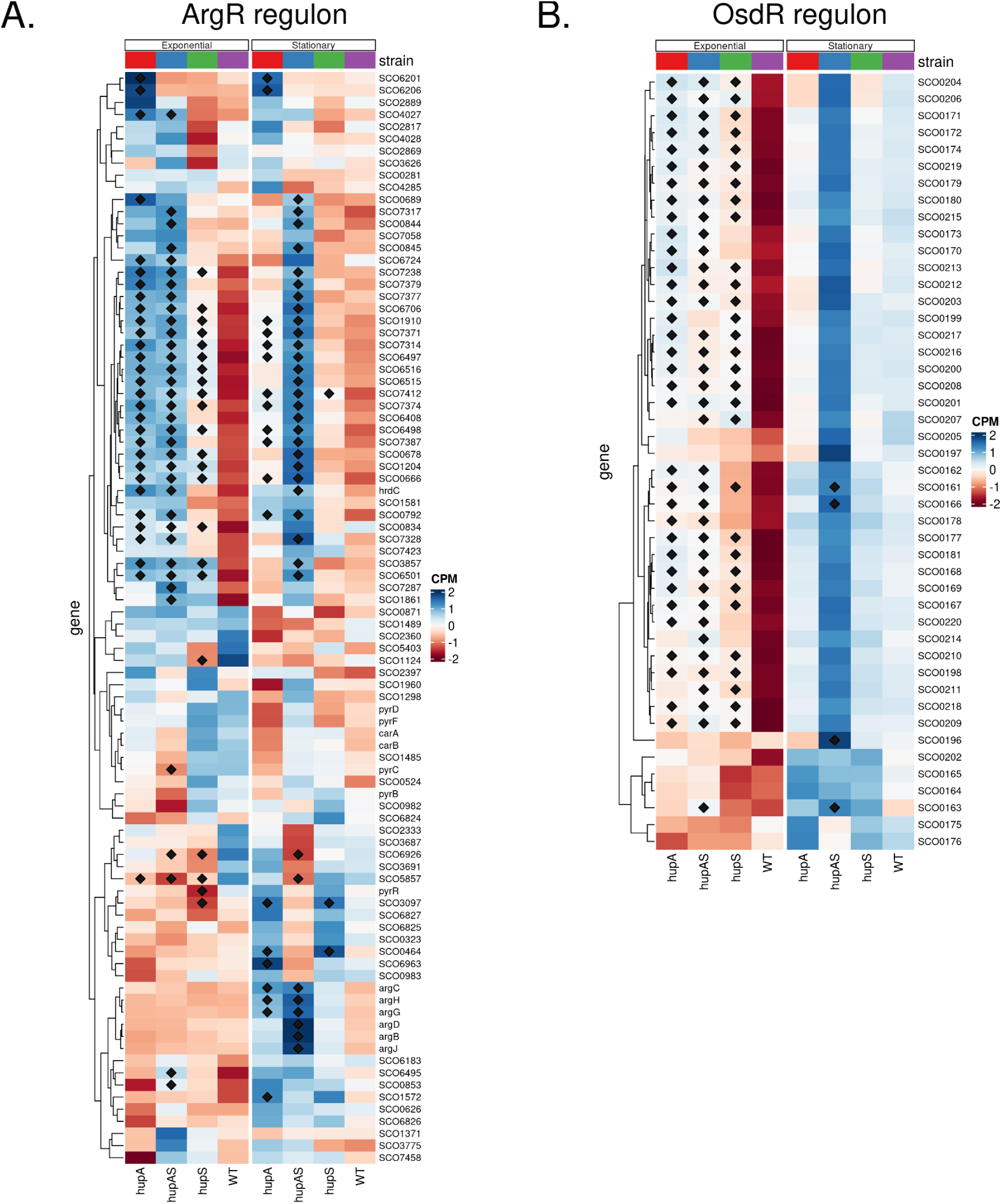
Genes clusters regulated by HupA and HupS. **A.** Heatmap showing normalized expression (CPM) of ArgR regulon genes for Δ*hupA,* Δ*hupS* and Δ*hupAΔhupS s*trains during exponential and stationary growth. **B.** Heatmap showing normalized expression (CPM) of OsdR regulon genes for Δ*hupA,* Δ*hupS* and Δ*hupA*Δ*hupS s*trains during exponential and stationary growth. **A, B** Genes marked with diamond were found to be significant in comparison to the wild type strain (FDR ≤ 0.05).

**Fig. S7.**
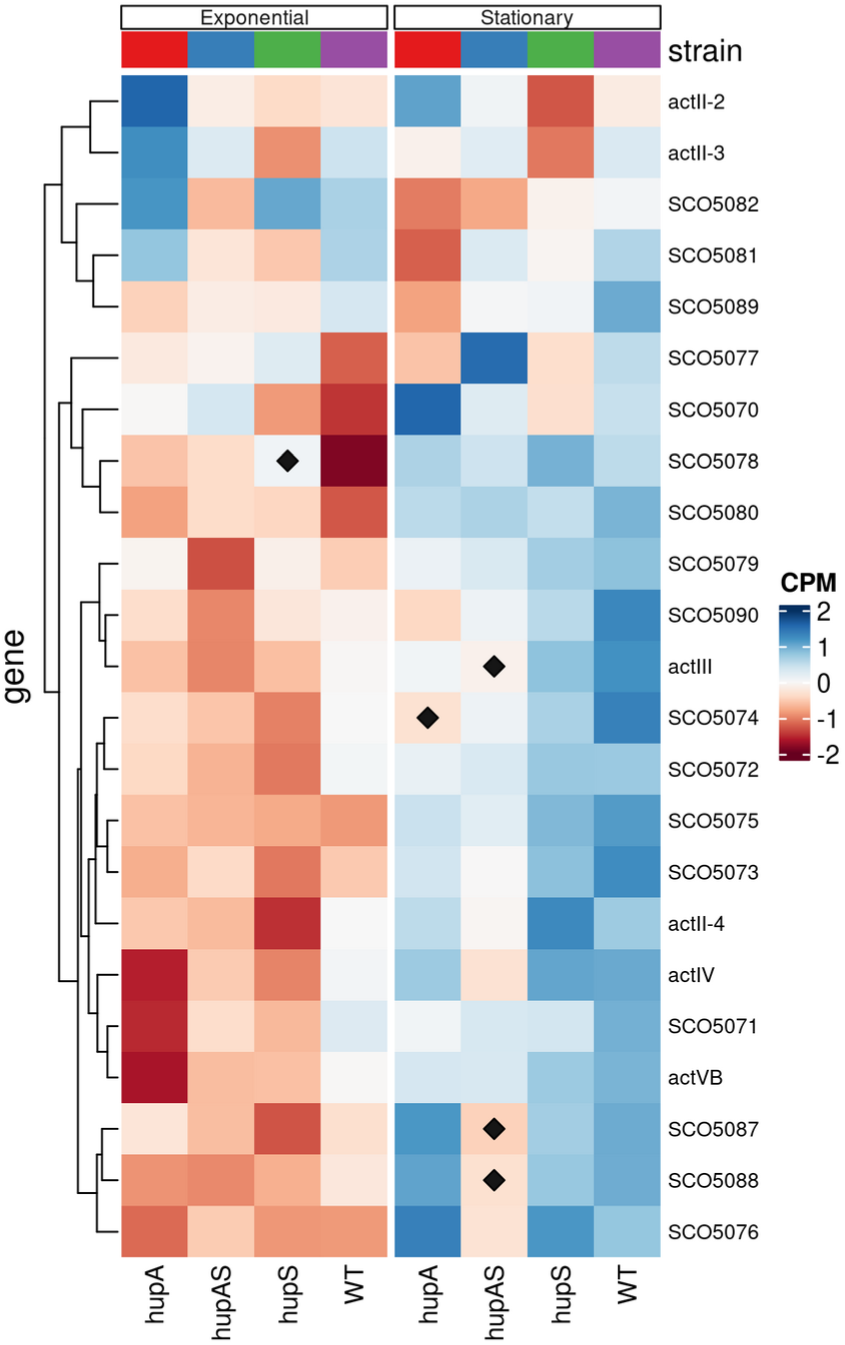
Heatmap showing normalized expression (CPM) of actinorhodin cluster genes for Δ*hupA,* Δ*hupS* and Δ*hupA*Δ*hupS* strains during exponential and stationary growth. Genes marked with diamond are those whose expression was found to be significantly altered in comparison to the wild type strain (FDR ≤ 0.05).

**Fig. S8.**
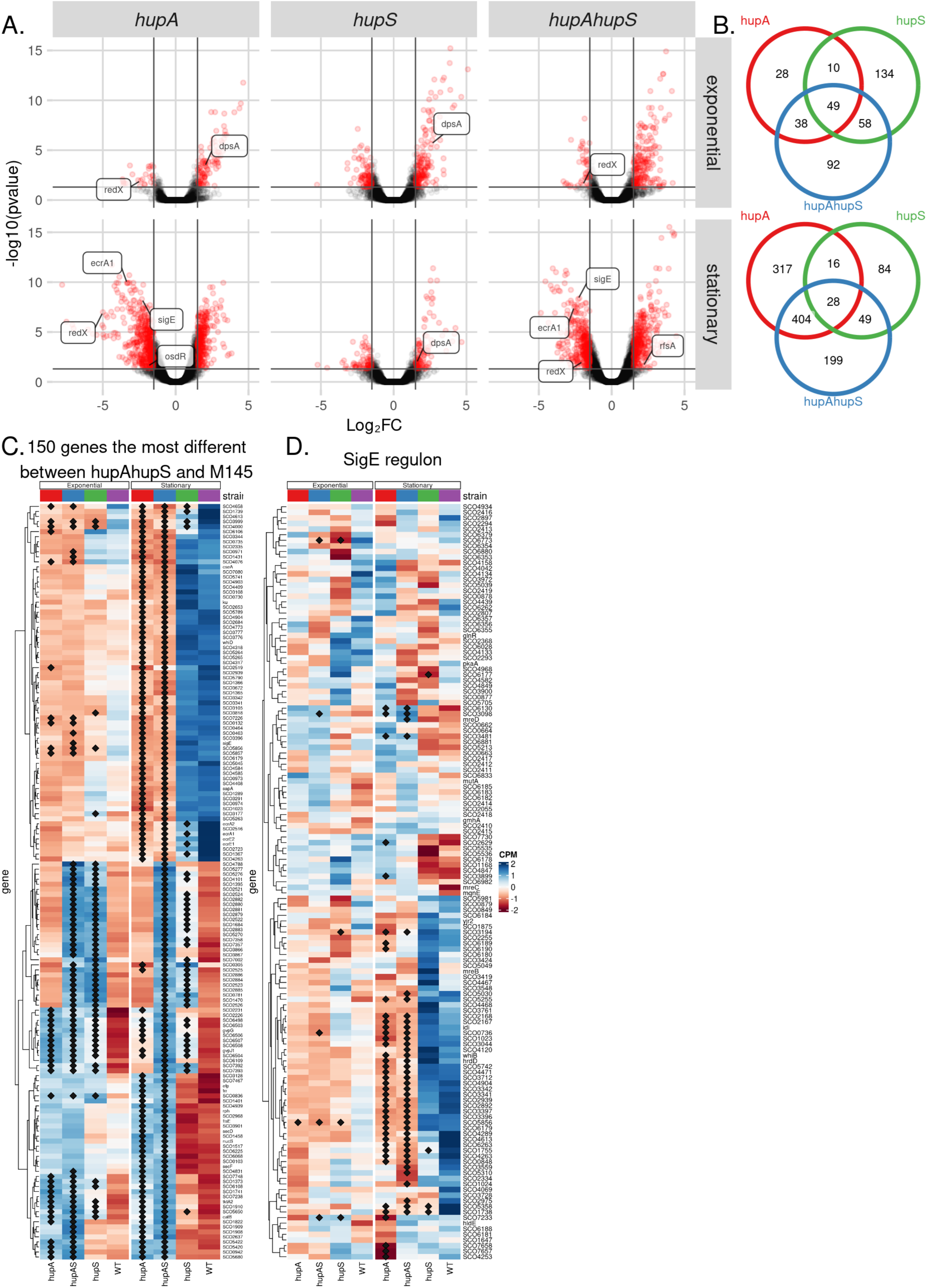
Changes of gene expression induced by osmotic stress conditions. **A.** Volcano plots showing modified gene expression in Δ*hupA,* Δ*hupS* and Δ*hupA*Δ*hupS* during exponential and stationary growth in medium supplemented with 0.5 M NaCl compared to the wild type strain. For each strain significantly changes genes (FDR ≤ 0.05, |Log_2_FC| > 1.5) are shown in red. Chosen differentially expressed genes are labelled. **B.** Venn diagrams showing significantly changed genes common for strains Δ*hupA,* Δ*hupS* and Δ*hupA*Δ*hupS* during exponential or stationary growth in medium supplemented with 0.5 M NaCl. **C.** Heatmap showing normalized expression (CPM) of 150 genes with the lowest FDR value calculated for Δ*hupA*Δ*hupS* strain and wild type comparison in wild type, Δ*hupA, ΔhupS* and Δ*hupAΔhupS s*trains during exponential and stationary growth in NaCl supplemented medium (0.5 M NaCl). **D.** Heatmap showing normalized expression (CPM) of SigE regulon genes for Δ*hupA, ΔhupS* and Δ*hupAΔhupS s*trains during exponential and stationary growth in NaCl supplemented medium (0.5 M NaCl). **C, D** Genes marked with diamond were found to be significant in comparison to the wild type strain (FDR ≤ 0.05, |Log_2_FC| > 1.5).

**Fig. S9.**
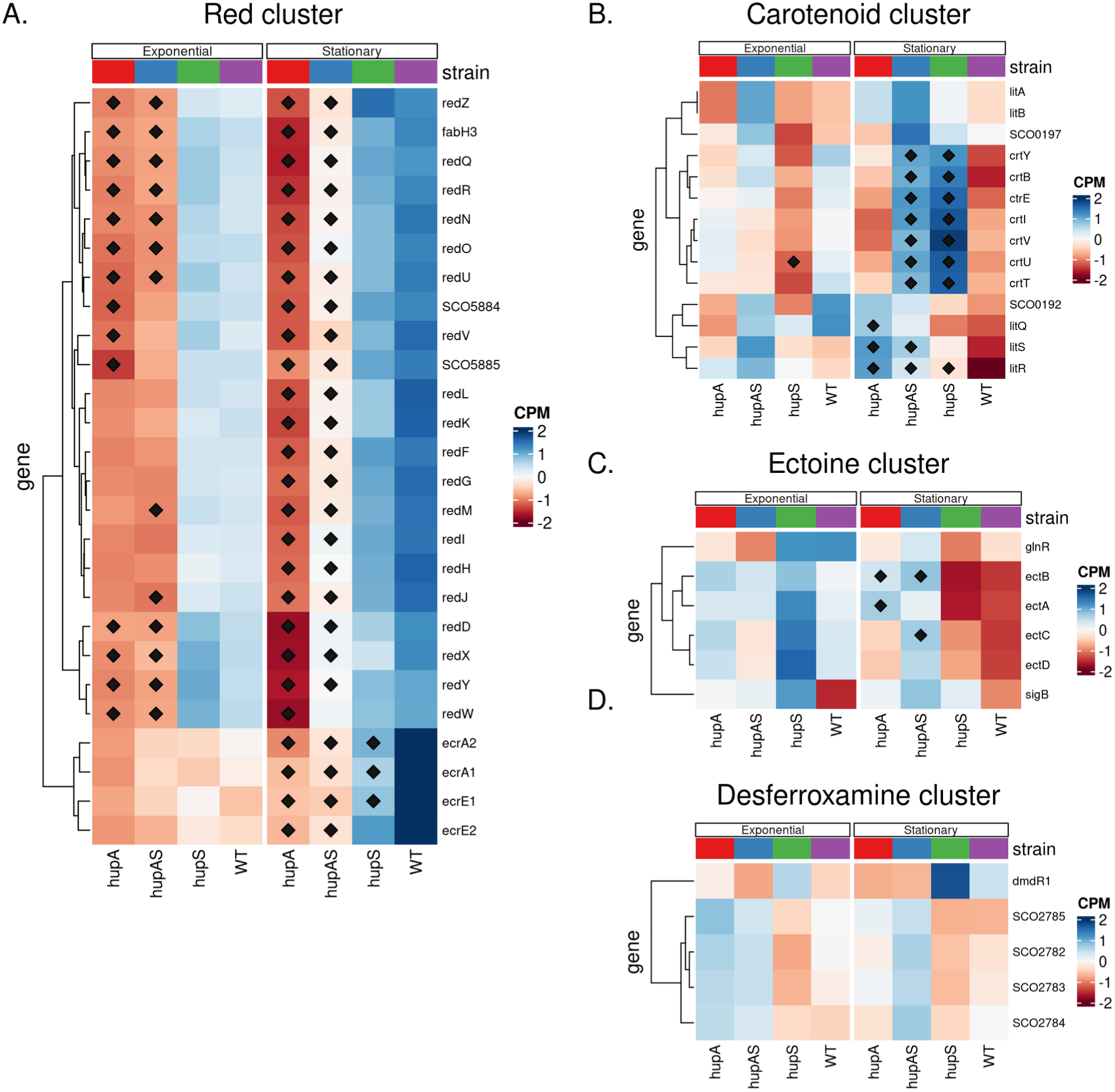
Modified expression of the genes within biostynthetic gene clusters, Δ*hupA,* Δ*hupS* and Δ*hupA*Δ*hupS* strains during growth at osmotic stress as compared to the wild type strain. **A.** Heatmap showing normalized expression (CPM) of the red cluster genes in wild type, Δ *hupA, ΔhupS* and Δ*hupA*Δ*hupS* strains during exponential and stationary growth in NaCl supplemented medium (0.5 M NaCl). **B.** Heatmap showing normalized expression (CPM) of the carotenoid cluster genes in wild type, Δ*hupA,* Δ*hupS* and Δ*hupA*Δ*hupS* strains during exponential and stationary growth in NaCl supplemented medium (0.5 M NaCl) **C.** Heatmap showing normalized expression (CPM) of ectoine cluster genes in wild type, Δ*hupA, ΔhupS* and Δ*hupAΔhupS s*trains during exponential and stationary growth in NaCl supplemented medium (0.5 M NaCl) **D.** Heatmap showing normalized expression (CPM) of desferroxamine cluster genes in wild type, Δ*hupA,* Δ*hupS* and Δ*hupA*Δ*hupS* strains during exponential and stationary growth in NaCl supplemented medium (0.5 M NaCl) **A, B, C, D** Genes marked with diamond were found to be significant in comparison to the wild type strain (FDR ≤ 0.05).

**Fig. S10.**
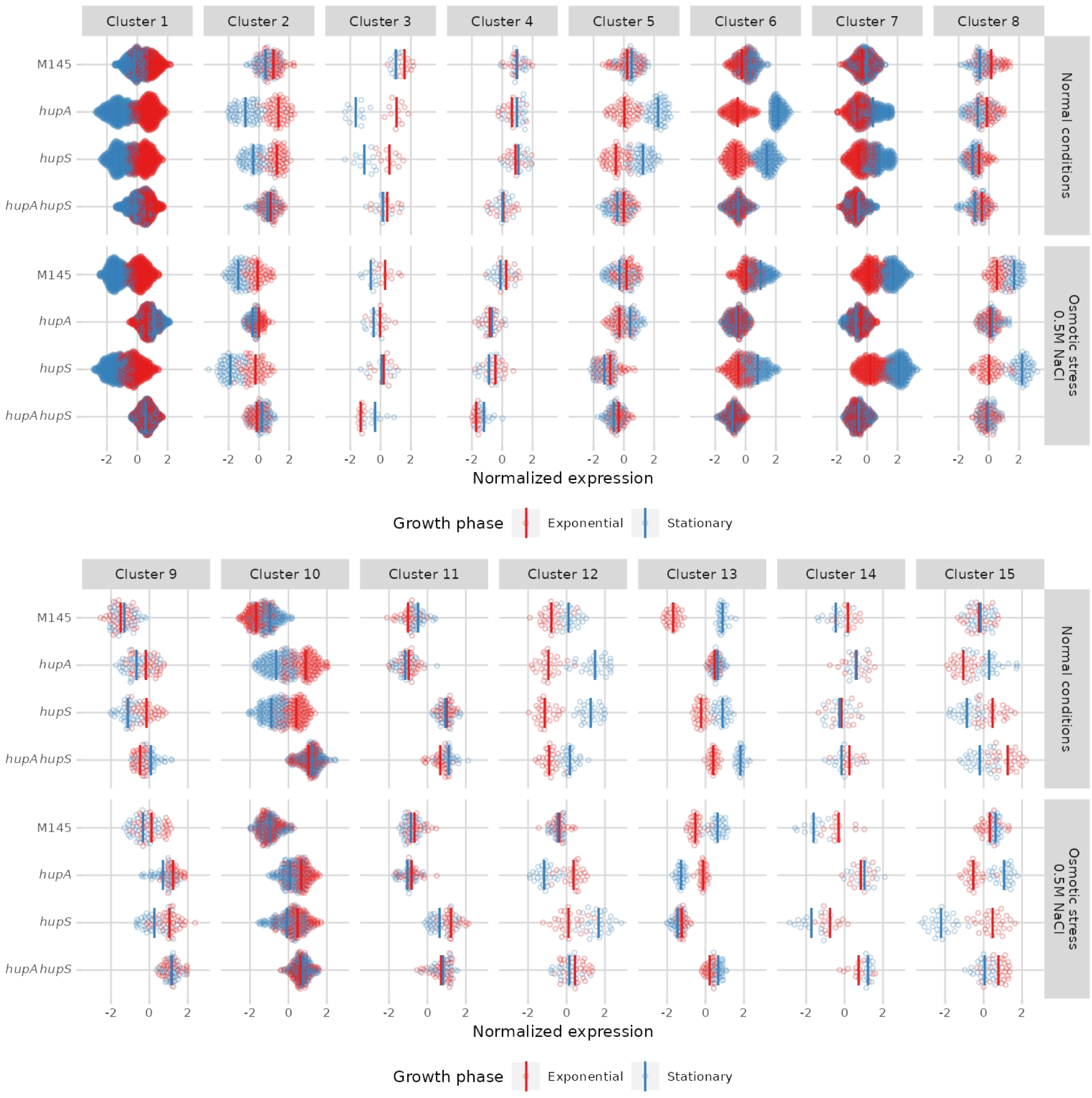
All clusters of genes identified based on transcriptional changes in Δ*hupA*, Δ*hupS* and Δ*hupA*Δ*hupS*. Results of clust analysis of wild type, Δ*hupA,* Δ*hupS* and Δ*hupA*Δ*hupS* transcriptomes from exponential (red) and stationary (blue) growth in normal and NaCl supplemented medium.

